# Fast Intensity Adaptation Enhances the Encoding of Sound in *Drosophila*

**DOI:** 10.1101/228213

**Authors:** Jan Clemens, Nofar Ozeri-Engelhard, Mala Murthy

## Abstract

To faithfully encode complex stimuli, sensory neurons should correct, via adaptation, for stimulus properties that corrupt pattern recognition. Here, we investigate sound intensity adaptation in the *Drosophila* auditory system, which is largely devoted to processing courtship song. Mechanosensory neurons (JONs) in the antenna are sensitive not only to sound-induced antennal vibrations, but also to wind or gravity, which affect the antenna’s mean position. Song pattern recognition therefore requires adaptation to antennal position (stimulus mean) in addition to sound intensity (stimulus variance). We discover fast variance adaptation in *Drosophila* JONs, which corrects for background noise over the behaviorally relevant intensity range. We determine where mean and variance adaptation arises and how they interact. A computational model explains our results using a sequence of subtractive and divisive adaptation modules, interleaved by rectification. These results lay the foundation for identifying the molecular and biophysical implementation of adaptation to the statistics of natural sensory stimuli.

## Introduction

Neural adaptation, the process of adjusting a neuron’s dynamic range to match the statistics of the sensory environment^1,2^, is ubiquitous throughout the nervous system and across species. Adaptation allows our visual systems to recognize a familiar face whether in the daytime or nighttime and our auditory systems to parse a sentence whether it is whispered or shouted. In general, adaptation corrects a neural representation for irrelevant aspects of the stimulus, such as fluctuations in stimulus mean or variance^2-4^. To ensure that the stimulus is efficiently encoded, adaptation should be both sufficiently fast and implemented as early as possible in the sensory pathway.

Accordingly, many sensory systems perform mean and variance adaptation within sensory receptor neurons. The nature of the physical stimulus transformed by receptor neurons into electrical activity dictates how they should implement adaptation. For auditory receptor neurons, the physical stimulus corresponds to the mechanical displacement of the receiver (for example, the stereocilia of cochlear hair cells). Receiver displacement has two, typically independent components: a fluctuating mean or baseline^5,6^ superimposed on sound-induced receiver oscillations, the standard deviation of which corresponds to the sound intensity. To faithfully encode the fine temporal features of sound, auditory receptor neurons should correct for both fluctuations in the mean and variance of receiver movements, without mean adaptation affecting sound sensitivity. Indeed, cochlear hair cells correct subtractively for stereocilia offsets (what we term here “mean adaptation”) and divisively for changes in intensity (what we term here “variance adaptation”)^7,8^, but how these forms of adaptation interact within hair cells has not yet been characterized. Studying these forms of adaptation in the more accessible auditory system of *Drosophila melanogaster* presents an opportunity to investigate their implementation.

Flies use sound for communication during their mating ritual. During courtship males chase females and produce dynamically-patterned song in bouts – females evaluate this acoustic signal and ultimately arbitrate the mating decision^9,10^. Song bouts typically comprise two modes – “sine” and “pulse”. To detect conspecifics and assess mate quality, the female’s auditory system must reliably encode the spectral and temporal properties of song, but several idiosyncrasies of both fly ears and fly courtship behavior can interfere with a faithful representation of song. The courtship song vibrates the arista (a feathery extension of the *Drosophila* antenna that serves as the sound receiver) – this causes rotation of the antenna and thereby opens mechanosensitive channels housed within antennal neurons, so-called Johnston’s organ (JO) neurons (JONs)^11,12^ (Fig. 1a, top). The male and the female constantly move during the courtship interaction^10^ and this induces strong fluctuations in the variance of sound-induced antennal movement (corresponding to the sound intensity, Fig. 1a bottom) for two reasons: i) fly ears are sensitive to the particle velocity component of sound (measured in mm s^-1^), which falls off strongly as the cube of the distance^13^, and ii) fly ears are sensitive to sound direction since the particle velocity is a directional quantity and the sound receiver is maximally vibrated only by sound coming from a direction orthogonal to the arista^14^. While males dynamically adjust how loudly they sing based on their distance to the female^15^, this adaptation on the sender side is likely insufficient to avoid saturation of the auditory receptors during courtship. The reported dynamic range of *Drosophila* auditory receptor neurons is between 0.1 and 1 mm s^-116^, but song can be as loud as 4 mm s^-1^ when the male is close to the female ^17^. In addition to intensity fluctuations, the baseline position of the antenna constantly shifts because of changes in air flow and gravity^11,12^. This baseline corresponds to the mean of the distribution of antennal displacements (Fig. 1a, bottom) and can saturate the responses to sound since mechanotransduction works only over a limited displacement range^18^.

**Figure 1:**
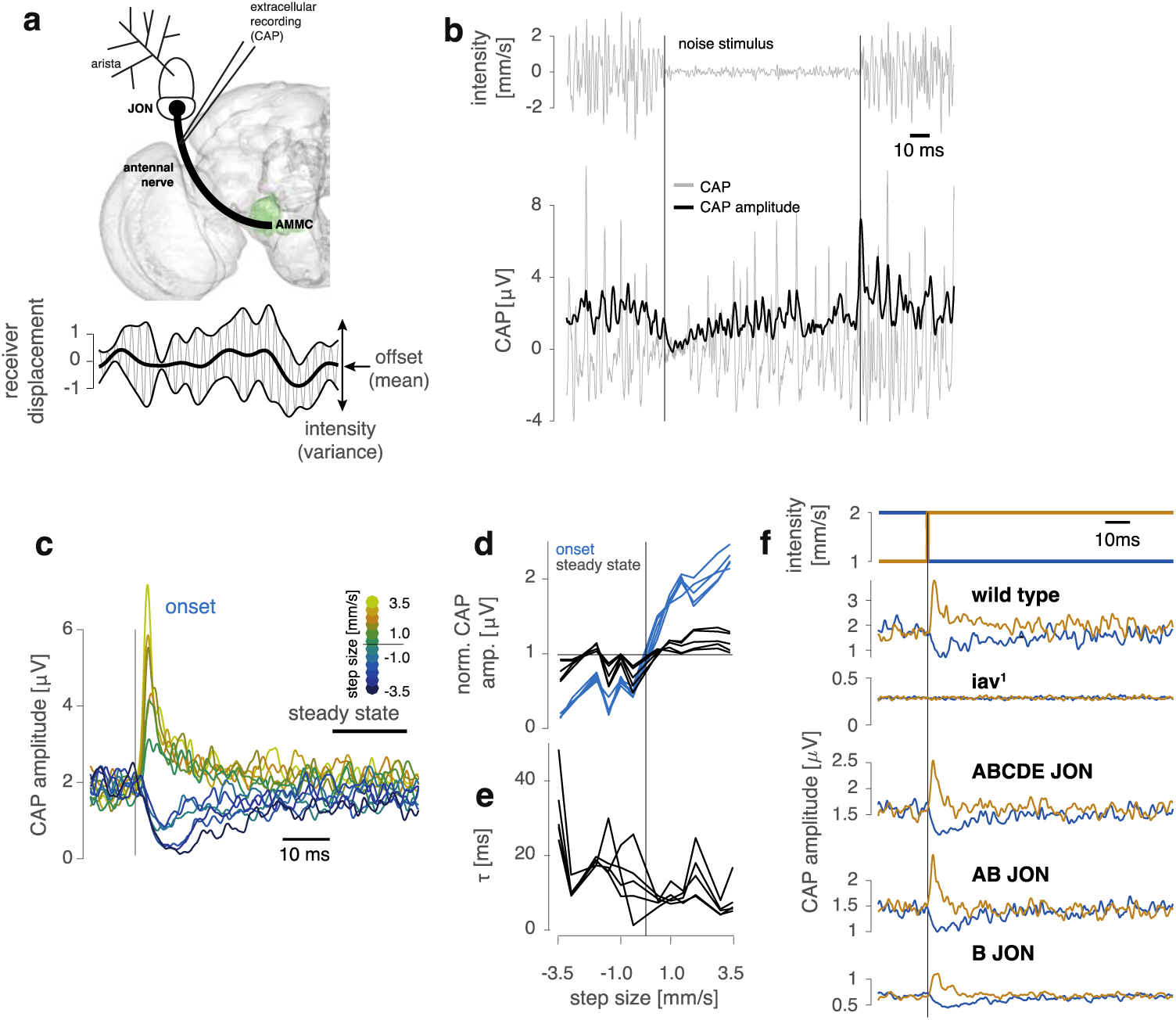
Sound intensity (variance) adaptation in JO neurons. **a** (top) Frontal view of the fly brain (left half) with schematic of the antenna. Sound-induced antennal vibrations activate antennal mechanosensitive neurons within the Johnston’s organ (JO) – a subpopulation of JO neurons (JONs) serve as the auditory receptor neurons. JON spikes travel down the antennal nerve to the antennal mechanosensory and motor center (AMMC, green) for central processing of acoustic stimuli. The compound spiking activity (CAP or compound action potential) of the JONs can be recorded extracellularly from the antennal nerve. (bottom) The mean (thick black line) of antennal displacements corresponds to a slowly changing offset; the variance (thin black lines) to the magnitude of sound-induced vibrations (sound intensity). **b** CAP recording (bottom, gray, averaged over 20 trials, see Supplementary Fig. 1b for single trial CAPs) for a noise stimulus (top) switching intensity every 100 ms. The CAP amplitude (bottom, black) was calculated using the Hilbert transform and exhibits transients after each intensity switch (vertical black lines). **c** CAP amplitude traces for all step sizes (color coded) aligned to step onset (vertical black line), for one fly. While the onset response strongly depends on step sign and magnitude, the steady-state response is relatively independent of step size. **d** Peak of onset and steady-state responses (blue and black, respectively) as a function of step size (N=5 flies). Normalized for each fly such that the average steady-state response across step sizes is 1.0. **e** Time scale of adaptation τ obtained by a fitting an exponential to the transients in the CAP amplitude (N=5 flies). **f** Responses of wild type, *iav^1^* mutants and *Iav*-rescues in different JON subsets to steps in noise intensity (from 1 to 2 mm s^-1^ (brown) and from 2 to 1 mm s^-1^ (blue), data from one fly each, see also Supplementary Fig. 1d-e). CAP signals are abolished in *iav^1^* flies but can be rescued by expressing functional *Iav* in specific JON subsets. CAP transients for all rescues resemble those seen in wild type. All CAPs averaged over 20 trials. Brain surface model (a) from https://github.com/jefferislab/nat.flybrains.

Thus, the *Drosophila* auditory system requires a neuronal mechanism for adaptation to both fluctuations in the baseline of antennal displacements and the intensity of the acoustic signal. While a subtractive mean adaptation mechanism corrects for the displacement baseline^18,19^, no studies have yet examined variance adaptation in *Drosophila*. Here, using neural recordings combined with computational modeling, we find that mean and variance adaptation co-occur within JONs and both do not require spiking or synaptic transmission. Despite the compact implementation of these two forms of adaptation within JONs, we find that mean adaptation does not affect sound sensitivity. This separation is not achieved by a separation of time scales since both forms of adaptation are equally fast. Rather, using a computational model, we show that the separation of mean and variance adaptation can be achieved with a simple sequence of computations: first, subtractive adaptation to correct for antennal offset, followed by rectification for encoding sound intensity, and finally divisive adaptation to correct for sound intensity. We find that the placement of rectification after mean adaptation in the model is essential for maintaining sound sensitivity for different levels of mean adaptation. While mean adaptation first occurs during mechanotransduction itself^18,19^, our data suggest that rectification and variance adaptation arise in subsequent steps. Our analysis of the organization of adaptation in *Drosophila* auditory receptor neurons now opens the door to employ genetic tools to identify the biophysical and molecular bases of these conserved operations, given the extensive parallels between fly and vertebrate hearing^18,20-22^.

## Results

### Variance adaptation in Drosophila auditory receptor neurons

The *Drosophila* sound receiver, its arista, is moved by sound and by slower, quasi-static stimuli, such as gravity or wind – these stimuli push and pull on the antenna, activating mechanosensory neurons (Johnston’s organ neurons or JONs) in its second segment (Fig. 1a). Antennal movement is actively amplified and produces a transduction or generator current in JON dendrites^18,19^. These currents are converted into action potentials that propagate along the antennal nerve to the antennal mechanosensory and motor center (AMMC), the first relay for auditory processing in the fly brain ^11^. While type CE JONs encode slow antennal movements by responding tonically to static displacement of the antenna^11,12,23^, type AB JONs rapidly and completely adapt their responses to these static displacements and produce sustained responses primarily for fast, sound-induced antennal vibrations in the frequency range of courtship song (100-350 Hz^18^. They hence act as the auditory receptor neurons^11,12^. However, it is unknown whether type AB JONs also adapt to sound intensity, corresponding to the variance of the sound-induced antennal vibrations superimposed on the quasi-static deflections caused by wind or gravity.

We first examined neural responses to band-limited Gaussian white noise stimuli (termed “noise” from here on) of fluctuating intensities. Since recording intracellularly from individual JONs with patch electrodes interferes with the natural movement of the antenna and with mechanotransduction (Supplementary Fig. 1a), we first probed for variance adaptation by recording population activity extracellularly from the antennal nerve (Fig. 1a). The compound-action potential (CAP) reflects the bulk spiking activity of sound-responsive JONs with sub-millisecond resolution (Supplementary Fig. 1b)^11,24^, and recordings are stable over hours. We presented a noise stimulus at intensities ranging from 0.25 to 2 mm s^-1^, with switches in intensity every 100 ms (Fig. 1b). Sound intensity is measured as particle velocity in mm s^-1^, since in the near field, bulk air movement, not changes in sound pressure, induce vibrations of the receiver (for *Drosophila* courtship, communication distance is much less than the wavelength of sound). The chosen intensity range corresponds to that which females naturally hear during courtship^17^. Noise exhibits a richly-patterned fine structure that evokes reproducible, temporally structured responses in the JONs and has been shown to reduce desynchronization in other systems^25^ (Supplementary Fig.1b, 2a).

We found that CAP responses exhibit transients at each switch in noise intensity: decreases in intensity produce negative transients while increases in intensity produce positive transients (Fig. 1c). After a few milliseconds, the responses settle to a steady-state that is relatively independent of sound intensity suggesting the presence of variance adaptation. To quantify these dynamic changes in sensitivity, we compared CAP amplitudes at the onset of an intensity switch and at steady-state – we did this for all intensity steps tested and for different instantiations of the white noise stimulus (see Methods for details). The resulting tuning curves reveal that the CAP amplitude strongly scales with step-size at the time of a switch but is nearly intensity-invariant at steady-state (Fig. 1d). Adaptation is fast, on the order of 5 to 20 ms, and its speed depends on the size of the intensity step, with positive steps producing the fastest and negative steps producing the slowest transients (Fig. 1e). Adaptation dynamics are asymmetrical likely because the amplitude of negative transients is limited from below since CAP amplitudes and the firing rates they represent can’t decrease to below 0Hz.

Since our extracellular recordings represent JON population activity, we ran several control experiments to determine that adaptive CAP dynamics specifically reflect adaptation of the firing rate within soundsensitive JO-AB neurons. First, we recorded the CAP in flies carrying the *iav^1^* mutation, which disrupts mechanotransduction and spiking in all JONs, and then rescued *iav* either in all JON, type AB or type B JONs^19,26^ (Supplementary Fig. 1d). The adaptation dynamics in these rescues are virtually indistinguishable from those of wild type flies (Fig. 1f, Supplementary 1e), suggesting the recorded CAP is dominated by activity from type AB JONs and that adaptation is a property of this subpopulation. This result is consistent with previous reports that show that the activity of type AB JONs, but not of type CE JONs, is visible in the CAP for the frequency and intensity range of stimuli used in our study^11,21,24^. Second, extracellular signals, such as cortical LFPs, can be sensitive to the synchrony among neurons in the population^27^. The noise stimuli we used should avoid desynchronization^25^, as can be shown using a simple model of a population of leaky integrate and fire (LIF) neurons (Supplementary Fig. 2). Only adding a positive offset to the stimulus would drive LIF neurons in a way that induces desynchronization, but this alone is not sufficient to produce transient changes in synchrony upon steps in noise intensity (Supplementary Fig. 3), further supporting the idea that desynchronization does not explain the CAP dynamics. Finally, we recorded calcium signals via GCaMP6f^28^ from type A and B JON projections into the AMMC (Supplementary Fig. 4a). Ca responses disappear when blocking JON spiking through bath application of TTX (Supplementary Fig. 4b), indicating that they represent the spiking activity of JONs. While the temporal resolution of GCaMP6f is impoverished compared with the CAP, calcium signals are not known to be sensitive to fine-scale neuronal synchrony and thus these recordings provide a more direct, albeit slow, readout of JON spiking. The calcium responses of both type A and B JONs for noise stimuli confirm the results obtained from CAP recordings (Supplementary Fig. 4c, d): responses saturate in the range of intensities tested (see also^16^) and exhibit both positive and negative transients immediately after steps. While adaptation appears to be weaker than that measured with the CAP likely due to the slow dynamics of the calcium indicator (Supplementary Fig. 4e), it is present nonetheless.

We also built simple, proof-of-principle models to demonstrate that alternative explanations of the adaptation dynamics in the CAP based on response heterogeneity – differences in frequency or intensity tuning – within type AB JONs are inconsistent with our data, in particular with the negative transients observed for negative steps in intensity (see Supplementary Fig. 5, 6 for details). Nonetheless, in the absence of recordings from individual type AB JONs, these models cannot rule out a contribution of desynchronization, intensity range fractionation, or other forms of response heterogeneity to our observation of variance adaptation. However, our models show that doing so will require auxiliary assumptions, and that changes in firing rate within individual type AB JONs is a more parsimonious explanation of all of our data.

The above experiments and analyses provide strong evidence that type AB JONs adapt their firing rates to changes in sound intensity (variance), but by what arithmetic operation do they do so? Since sound intensity scales the distribution of antennal displacements, variance adaptation should be divisive, not subtractive, and result in a change of the slope of the intensity tuning curve. We therefore measured JON responses to short, intermittent probe stimuli interleaved within background noise stimuli of a particular intensity (Fig. 2a, bottom). The resulting family of intensity tuning curves – one for each background intensity – describes how JON sensitivity changes for different adaptation states (Fig. 2a, top). On a linear intensity scale, the slope of the tuning curves decreases with background intensity, indicating that variance adaptation is indeed divisive. A logarithmic intensity scale, which transforms this division to a rightward shift of the tuning curves, reveals a) that tuning curve shape is invariant to intensity (Fig. 2b, c) and b) that the shift is proportional to the background intensity (Fig. 2d). Variance adaptation in JON thus completely corrects for intensity across the dynamic range tested – in both male and female flies (Supplementary Fig. 7). Regardless of the shift, the tuning curve’s steepest part is always centered around the background level. Accordingly, stimulus discriminability – quantified using Fisher information^29^ – also shifts dynamically and is always maximal around the background level (Fig. 2e-g). Thus, adaptation enhances the resolution with which changes from the background level are encoded. More broadly, JONs do not encode absolute stimulus intensity, but intensity *relative* to the background, as is the case for many other sensory systems^30,31^.

**Figure 2:**
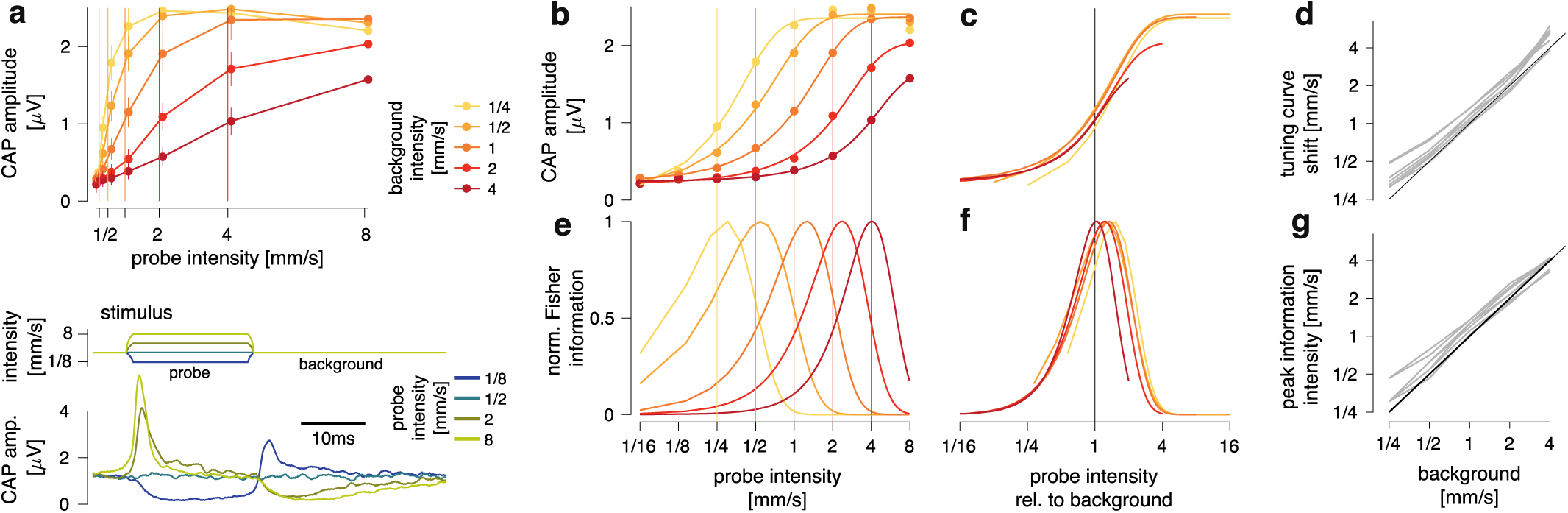
Variance adaptation in JON Is divisive and corrects for background noise. **a** Intensity tuning curves for JONs (data from one fly) adapted to different background stimuli (top, color coded by background intensity; vertical lines mark the background intensity; 40 trials for each background and probe combination). JONs were adapted using a white noise background sound and sound sensitivity was measured using brief, interleaved probe stimuli (amplitude profile of the noise stimulus shown in bottom). On a linear intensity scale, the slope of the tuning curves decreases with background – variance adaptation is thus divisive. Error bars show mean ± 95% CI. **b** Same as a but plotted on a logarithmic intensity scale, which transforms the change in slope into a tuning curve shift. Dots correspond to data (error bars omitted for clarity), lines show fit of sigmoidal curves (see Methods for details). **c** Sigmoidal fits from b plotted on a log-intensity scale relative to background intensity. Tuning curves overlap, demonstrating that the tuning curve shift is proportional to the background intensity. d Shift of the tuning curves is proportional to background intensity for all N=5 flies (r^2^=0.95). The diagonal line corresponds to a perfect match between the tuning curve position and background. All JON tuning curves largely match the diagonal’s slope with a weak positive offset. **e, f** Fisher Information curves for the tuning curves in b (normalized to max) show that the discriminability of intensities shifts in proportion to the background intensity (e – absolute log intensity, f – backgroundrelative log intensity). **g** The peak intensities for the Fisher Information curves shift with background for all N=5 flies (r^2^=0.97).

### Variance adaptation for song stimuli

We next examined the properties of variance adaptation for naturalistic stimuli. *Drosophila* courtship song comprises bouts consisting of rapid alternations between two modes – sine and pulse (Fig. 3a). Sound can be decomposed into two components: fast fluctuations of the raw waveform (the carrier), and an envelope, which corresponds to amplitude changes of the carrier on a slower timescale (Fig. 3b). Sine song and pulse song have different, species-specific, sinusoidal carriers of ~150 and 250-300 Hz, respectively^32,33^. But song modes also differ in their envelope dynamics: pulse song’s envelope is highly transient, with a species-specific interval between pulses in a pulse train (inter-pulse interval) of ~36ms for *melanogaster*. In contrast, the envelope of sine song is only weakly modulated and hence resembles the steady envelope of noise stimuli (Fig. 3b).

**Figure 3:**
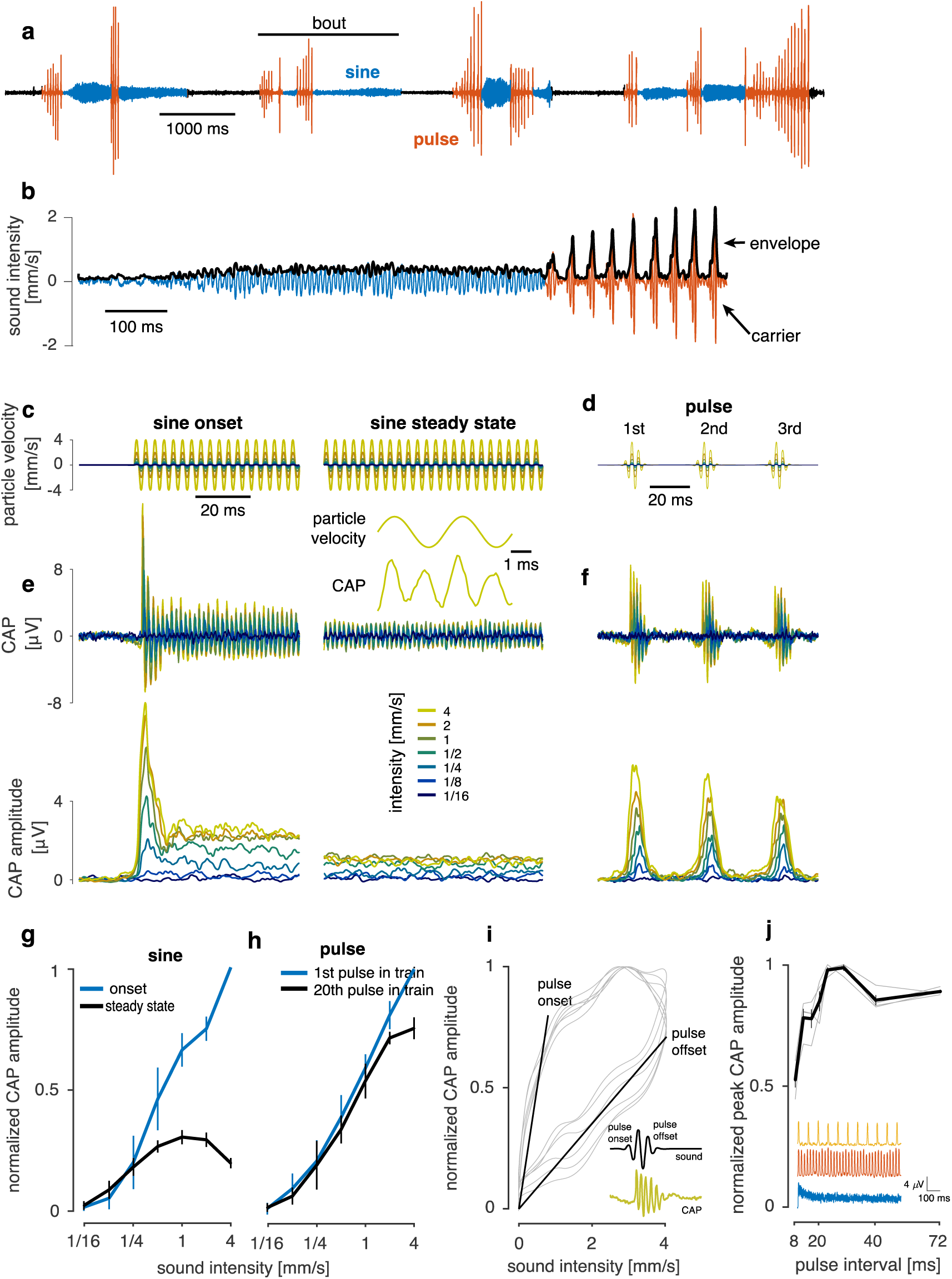
Variance adaptation and courtship song signals. **a** Courtship song is produced in bouts and consists of two modes: sine (blue) and pulse (red). **b** Sine song (blue) exhibits a relatively constant amplitude profile, while pulse song (red) consist of trains of transient pulses with interleaved pauses. **c, d** Artificial sine song and pulse song presented at a range of different intensities (color coded, see legend in e). **e** CAP (top – raw CAP, bottom – CAP amplitude) from one fly for 300 Hz sinusoids of different intensities (c). Intensity is color coded (see legend). After a strong onset transient (left), the CAP amplitude drops rapidly (right – steady-state response after 500ms). Inset shows the frequency doubling of the CAP (top – stimulus at 4 mm s^-1^, bottom – CAP response oscillates at twice the stimulus frequency, or 600Hz). **f** CAP (top – raw CAP, bottom – CAP amplitude) from one fly for synthetic pulse trains with different intensities (d, pulse interval 36 ms, intensity color-coded, see legend in e). While there is a weak decrease of response amplitude across pulses in a train, responses still scale with intensity (see also Supplementary Fig. 8c-e). **g, h** Onset and steady-state intensity tuning for sine stimuli (g) and pulse stimuli (h) (see also Supplementary Fig. 8); N=6 flies, 20 trials each. Error bars correspond to mean ± 95% CI across flies. **i** Asymmetrical responses to pulses (inset – CAP from one fly) betray adaptation. Thin black lines show CAP amplitude vs. stimulus amplitude for the first six pulses in a train (pulse peak intensity 2 mm s^-1^). N=6 flies. Responses to pulse onset have a much larger slope then those to pulse offset (see diagonal lines). **j** Peak responses at steady-state (after 1.4 seconds into pulse train) for pulses trains with different interpulse intervals (pulse duration 16ms, intensity 4mm s^-1^, gray lines show response of individual flies (N=4), normalized to the max response; thick black lines correspond to mean±s.e.m.). Responses are weak for short inter-pulse intervals due to stronger adaptation. Inset shows CAP amplitude traces for pulse trains with IPIs of 8ms (blue), 24ms (red) and 72ms (orange). All CAP traces averaged over 20 trials.

We examined neural responses for song-like stimuli across a range of intensities (0.06-4mm s^-1^, Fig. 3c-d). Consistent with our expectation, adaptation to sinusoidals resembles that of adaptation to noise (compare with Fig. 1b-c), with onset transients that scale with intensity and weaker, saturating steady-state responses (Fig. 3e, g, compare with Fig. 1d). This adaptation is not confined to the *melanogaster* sine song frequencies but occurs at all frequencies tested (from 100-900 Hz, Supplementary Fig. 8a, b). Adaptation was much weaker for pulse song (Fig. 3d), and intensity tuning resembled that of sine song *onsets* throughout the pulse train, with a near-linear relation between intensity and response (Fig. 3f, h). Nonetheless, other features of the responses to pulse trains betrayed a dynamic adjustment of response gain. First, pulse response amplitude weakly decreased across pulses in a train particularly at high intensities (Supplementary Fig. 8c-e). Second, responses to symmetrical pulses were asymmetrical, with strong responses during pulse onset and shallower responses during pulse offset (Fig. 3i). Asymmetry of responses for on and off ramps is a signature of gain control in other systems, such as the olfactory receptor neurons of *Drosophila* larvae^34^. Additionally, when we tested stimuli of different inter-pulse intervals, we found that the responses to individual pulses depended on the interval between pulses, with short intervals inducing weaker responses during the pulse train, due to stronger adaptation (Fig. 3j). We also observed a small reduction of responses for the longest IPIs tested, which cannot be explained by adaptation: CAP responses recover from adaptation within ~30 ms and hence adaptation does not affect responses to pulses that are spaced further apart in time (Fig. 1d). Overall, although variance adaptation is too slow to correct for the amplitude of individual pulses at the *melanogaster*-typical pulse interval, it acts as a high-pass filter for this species-specific song feature (reducing responses to songs with shorter IPIs), and thereby could contribute to song evaluation.

### Cellular origins of variance adaptation in the antenna

Since there are no synapses between individual JONs (with the possible exception of gap junctions between JON axons^35^) nor feedback to the JON from the brain or other sensory structures^36^, variance adaptation is likely to arise within individual JONs, at one of the following levels (Fig. 4a): i) the movements of the antenna, ii) the subthreshold mechanosensitive generator currents^19,21^, or iii) the generation of spikes.

**Figure 4:**
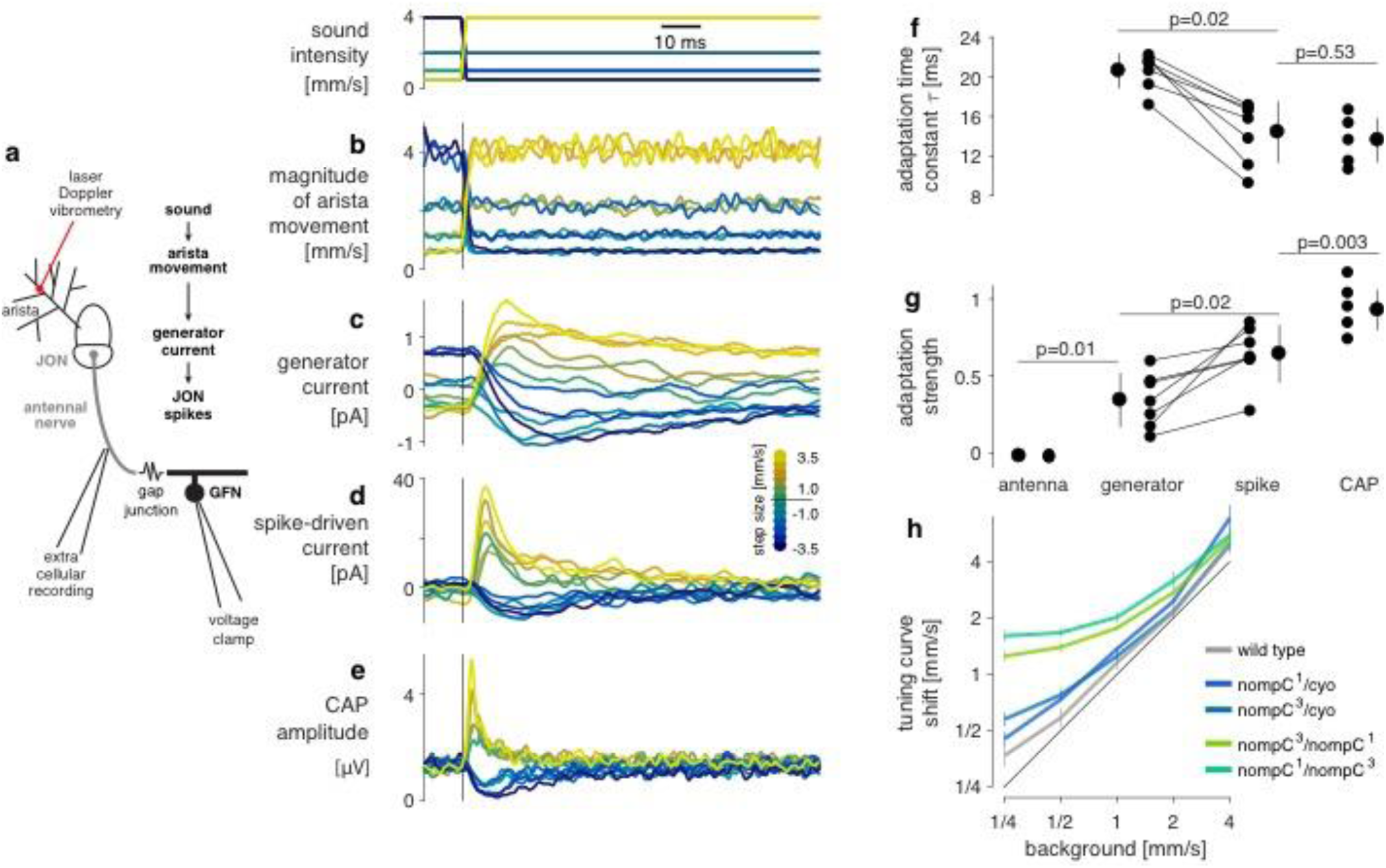
Cellular basis of variance adaptation in JO neurons. **a** Sound-induced antennal movement was measured using laser vibrometry (b). Generator (c) and spike-driven (d) currents were read out via patch clamp recordings of the giant fiber neuron (GFN), after bath application of TTX (c) or with normal saline (d). **b-e** Amplitude profile of the noise steps used to probe adaptation in different processing stages in JONs (top). Step size is color coded (see legend in d). Magnitude of antennal movement (b), generator current (c), spike-driven currents (d) and CAP (e) to the intensity steps. Adaptation transients first arise in the generator current (c). All panels show one fly with 20 trials per step (c – 100 trials per step). **f** Time constant of adaptation at different stages within JON processing obtained by fitting exponentials to the transients in b-e (time constant averaged across all step sizes). The lack of adaptation in the antennal movements prevented estimation of time constants for the antenna. P-values indicate the results of two-tailed tests (sign test for paired data (generator vs. spike), rank sum for unpaired data (LVM vs. generator, spike vs. CAP)). N = 7 flies for generator/spike, N=4 for LVM, N=5 for CAP (number of trials as in b-e). **g** Strength of adaptation for the signals in b-e was obtained by comparing the responses to the different intensities at step onset versus steady-state. Adaptation values of 1.0 indicate complete adaptation (values can be greater than 1 or smaller than 0 due to noise in the CAP responses). Statistical procedures and N as in f. Error bars in f and g show mean ± standard deviation over flies. **h** Adaptation in flies mutant for the TRP channel *nompC* (green, blue, wild type flies shown in gray for comparison). All mutants adapt for background intensity by shifting their tuning curve. The black diagonal line corresponds to perfect adaptation, complete absence of adaptation would lead to horizontal graphs. Adaptation is reduced for soft backgrounds (1/4-1mm s^-1^) in the *nompC* double mutants due to their reduced sound sensitivity. Mean ± s.t.d. over flies (see Supplementary Fig. 9 for N).

Like cochlear hair cells, the JO neurons are not passive transducers of mechanical stimuli but actively amplify antennal movement for soft sounds and partly reset antennal position after a static deflection through mean adaptation^16,18,21,22,37^. If the variance adaptation reported here was part of the same process underlying amplification and mean adaptation, then it should reduce the gain of antennal movement. This should decrease sound-induced antennal vibrations following a rapid increase in sound intensity and create adaptation transients in antennal position. We measured arista movement using laser Doppler vibrometry and found that the amplitude of antennal vibrations faithfully followed changes in stimulus intensity (Fig. 4b), lacking the adaptation transients seen in the CAP (Fig. 4e). We also tested *nompC* mutants that lack active amplification^37^. These mutants have diminished sound sensitivity^16^, but nonetheless we found that for stimuli that evoked strong responses (> 1 mm s^-1^), variance adaptation was still present (Fig. 4h and Supplementary Fig. 9). That is, JON intensity tuning shifted with background intensity in the *nompC* background, just as for wild type. These results suggest that variance adaptation in the JONs is independent of active antennal mechanics and amplification.

Following mechanical amplification, sound-induced antennal vibrations open stretch-sensitive ion channels in the JON’s dendritic tips and produce a transduction or generator current^19^. This subthreshold current could adapt either directly, as part of the mechanotransduction machinery, or indirectly, on its way to the spike-initiating zone (Fig. 4a). While the generator currents are too small to be recorded extracellularly, a subset of type A JONs form gap junction-only synapses with the giant fiber neuron (GFN)^38-40^ and recording the GFN in voltage-clamp mode reveals the synaptic currents induced by the spikes of type A JONs^19^. These spike-induced currents in the GFN provide an alternative readout of JON spiking activity and exhibit adaptation similar to that of the CAP, confirming that the CAP is an appropriate readout of JON spiking (Fig. 4d, compare with CAP in Fig. 4e). Bath-application of TTX during these GFN recordings abolishes the spike-driven input current from the JONs and unmasks the generator current^19^, which exhibits transients upon stimulation with noise steps, demonstrating that adaptation is present in the sub-threshold currents (Fig. 4c).

The adaptation transients recorded from the GFN are slower than those in the CAP (Fig. 4f) and adaptation appears less complete (Fig. 4g). This may be because the generator current travels passively along the axons of type A JONs and through the gap junction to the recording site at the GFN soma; this pathway likely acts as a low-pass filter. Alternatively, subsequent spike-frequency adaptation with JONs could further accelerate and complete variance adaptation. Nonetheless, the existence of transients demonstrates that variance adaptation arises in the generator current and does not require spiking. Because adaptation in many sensory neurons involves the calcium-sensitive molecule Calmodulin^7,41-43^, we also tested adaptation in *calmodulin* mutants^20^. However, these flies exhibited normal adaptation (Supplementary Fig. 9), suggesting that variance adaptation in JONs either operates via a calcium independent mechanism or another calcium-sensitive molecule^44^. So far we have shown that physiologically separable, computations in *Drosophila* auditory receptor neurons appear prior to spiking within individual JONs: active amplification^37^, mean adaptation^18,19,22^ and now variance adaptation. This raises the question of how these two forms of adaptation interact, given that sound sensitivity should be independent of the slow antennal offset induced by wind or gravity.

### Interaction between mean and variance adaptation

To explore the interaction between mean and variance adaptation, we controlled the antennal offset (or mean position of the antenna) and the magnitude (or variance) of antennal vibrations independently using a piezoelectric actuator. Piezoelectric actuation reproduced previous results regarding mean adaptation studied using electrostatic forces to deflect the antenna^18^ (Supplementary Fig. 10). We first engaged mean adaptation by statically deflecting the antenna (to +/-0.22 or +/-0.44 μm) and tested whether this affected tuning for intensity by probing with sinusoidal motion of different magnitudes (300 Hz, 0.1 to 1.1 mm s^-1^, Fig. 5a). We found that intensity tuning remained the same for all offsets, demonstrating that mean adaptation does not affect sensitivity to sound (Fig. 5b, c). On the other hand, when we engaged variance adaptation via sinusoidal modulations (300 Hz, 0.07 to 0.63 mm s^-1^) and subsequently measured responses to superimposed steps (+/- 0.01 to 0.78 μm, Fig. 5d), sensitivity to step stimuli decreased (Fig. 5e, f). Thus, the interaction between mean and variance adaptation in JONs is unidirectional – variance adaptation affects responses to displacement but mean adaptation does not affect responses to sound. This suggests that the majority of variance adaptation occurs downstream of mean adaptation, consistent with the fact that mean adaptation is already visible at the level of antennal movement^18^, but variance adaptation is not (Fig. 4b). The independence of mean and variance adaptation could be achieved through a separation of time scales, but we find that both forms of adaptation are similarly fast^18^ (Fig. 1e, Supplementary Fig. 10). To gain insight into the computations underlying the independence of sound sensitivity from mean adaptation, and to test the hypothesis of a serial implementation of first mean and then variance adaptation, we built a phenomenological model of adaptation in the JO.

**Figure 5:**
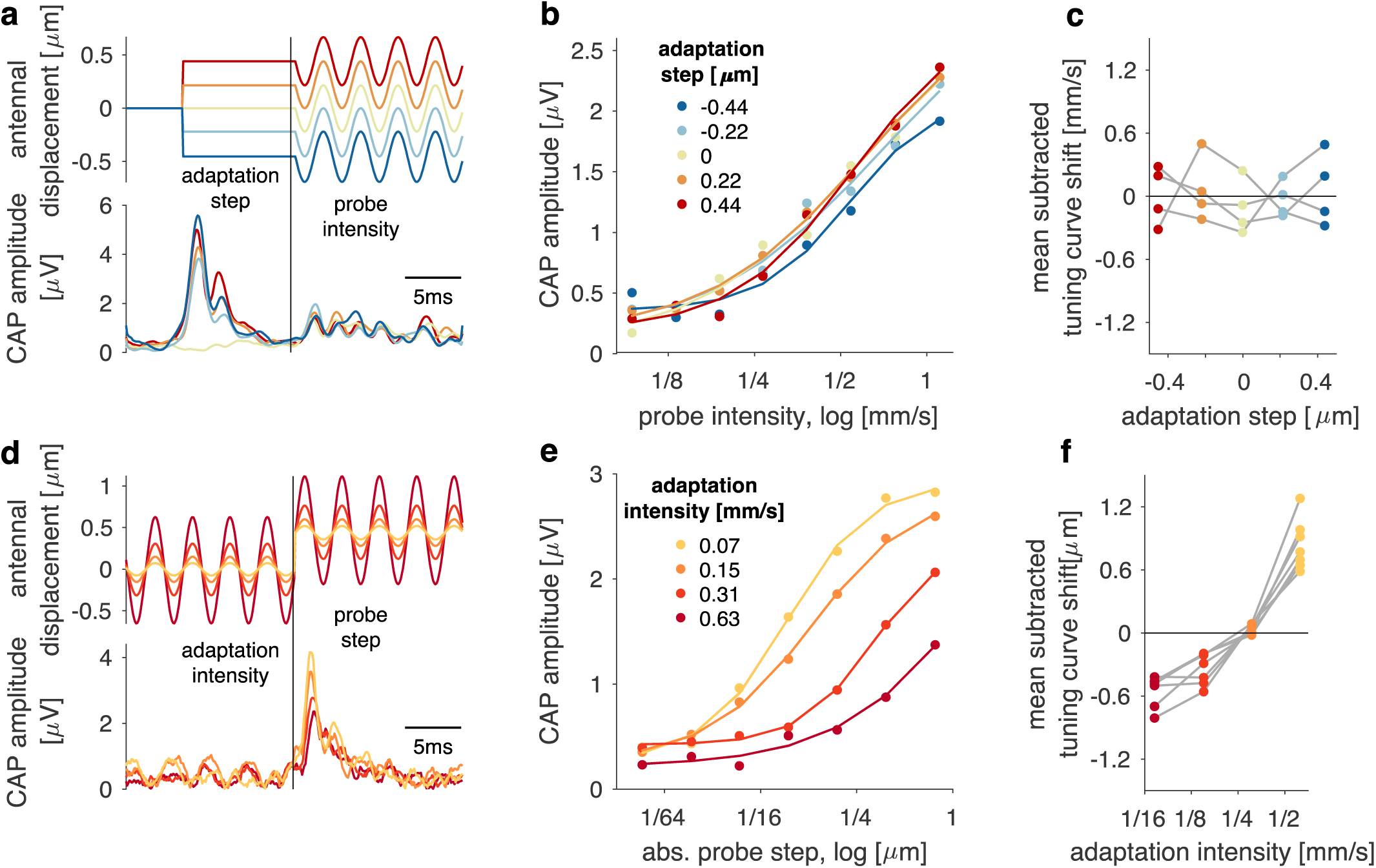
Interaction between offset (mean) and intensity (variance) adaptation in JO neurons. **a** Stimuli for probing the impact of offset (mean) adaptation on intensity tuning (top) – a piezoelectric actuator was used to step-deflect the antenna to induce mean adaptation (see Supplementary Fig. 10). Sensitivity to sound intensity was estimated by measuring the responses (bottom) to sinusoidal antennal deflection superimposed on the steps of different size (color coded – see insert in b). **b** Intensity tuning curves (dots) and sigmoidal fits (lines) for different levels of mean adaptation (color coded, see legend). **c** Tuning curve shifts with offset extracted from the sigmoidal fits (see b) for N=4 flies. No significant correlation with the adaptation step was observed for any fly (p>0.14). **d** Impact of variance adaptation on step responses was probed by superimposing step deflections onto sinusoidal antennal modulation of different magnitudes, again via piezoelectric actuator, and measuring responses (color coded - see insert in d). **e** Step tuning curves (dots) and sigmoidal fits (lines) for different levels of variance adaptation (color coded, see legend). Since step tuning curves are symmetrical, responses to positive and negative steps were averaged. **f** Tuning curve shifts with intensity extracted from the sigmoidal fits (see e) for N=7 flies. Significant correlation with adaptation step was observed in all flies (p<0.01). a, b, d, and e show responses from one fly, each averaged over 20 trials.

### A sequential network motif for mean and variance adaptation

We started by defining a set of elementary computations we deemed necessary for reproducing our experimental findings. Mean and variance adaptation in our model are implemented using incoherent feed-forward loops^34,45^: The loop’s input is low-pass filtered to an adaptation signal, which is either subtracted from the input (subtractive adaptation) or used to divide the input (divisive adaptation). We also included a rectifying nonlinearity in our model, since rectification is known to be necessary for variance adaptation^46^.

To explore the constraints underlying the organization of the two types of adaptation we consider here, we tested all possible sequential or parallel arrangements of these three elementary computations in our model (Supplementary Fig. 11a). Consistent with our hypothesis, only a sequential arrangement of 1) subtractive adaptation, 2) rectification and 3) divisive adaptation reproduced our experimental observations (Fig. 6a). This sequence corresponds to the solution engineers would choose if they wanted to correct a signal for its mean and variance: first subtract the mean, estimate the signal’s intensity through rectification and then divide by the intensity. The model’s qualitative behavior was robust to changes in model parameters (e.g., adaptation time constants or adaptation strength, Supplementary Fig. 11 b, c). In particular, the independence of sound sensitivity from mean adaptation does not require a separation of time scales – it persists even when mean adaptation is faster than variance adaptation (Supplementary Fig. 11b). The resulting pattern of adaptation is thus a general property of this arrangement of computations, not of the specific parameters chosen. The first stage of the network motif involves low-pass filtering the stimulus to produce an adaptation signal that represents the slowly-varying stimulus mean, which is then subtracted from the stimulus (Fig. 6b, c). For step stimuli, this adaptation signal corresponds to a smoothed version of the stimulus (Fig. 6b). For sound stimuli, however, with their symmetrical and fast fluctuations, this signal is nearly flat, since the positive and negative deflections cancel each other during the low-pass filtering (Fig. 6c). This first stage thus only adapts to quasi-static stimulus components (Fig. 6f, compare Supplementary Fig. 10) but does not alter responses to the sound (Fig. 6c). This is why mean adaptation does not affect sound sensitivity in the model (compare Fig. 6g with Fig. 5b).

**Figure 6:**
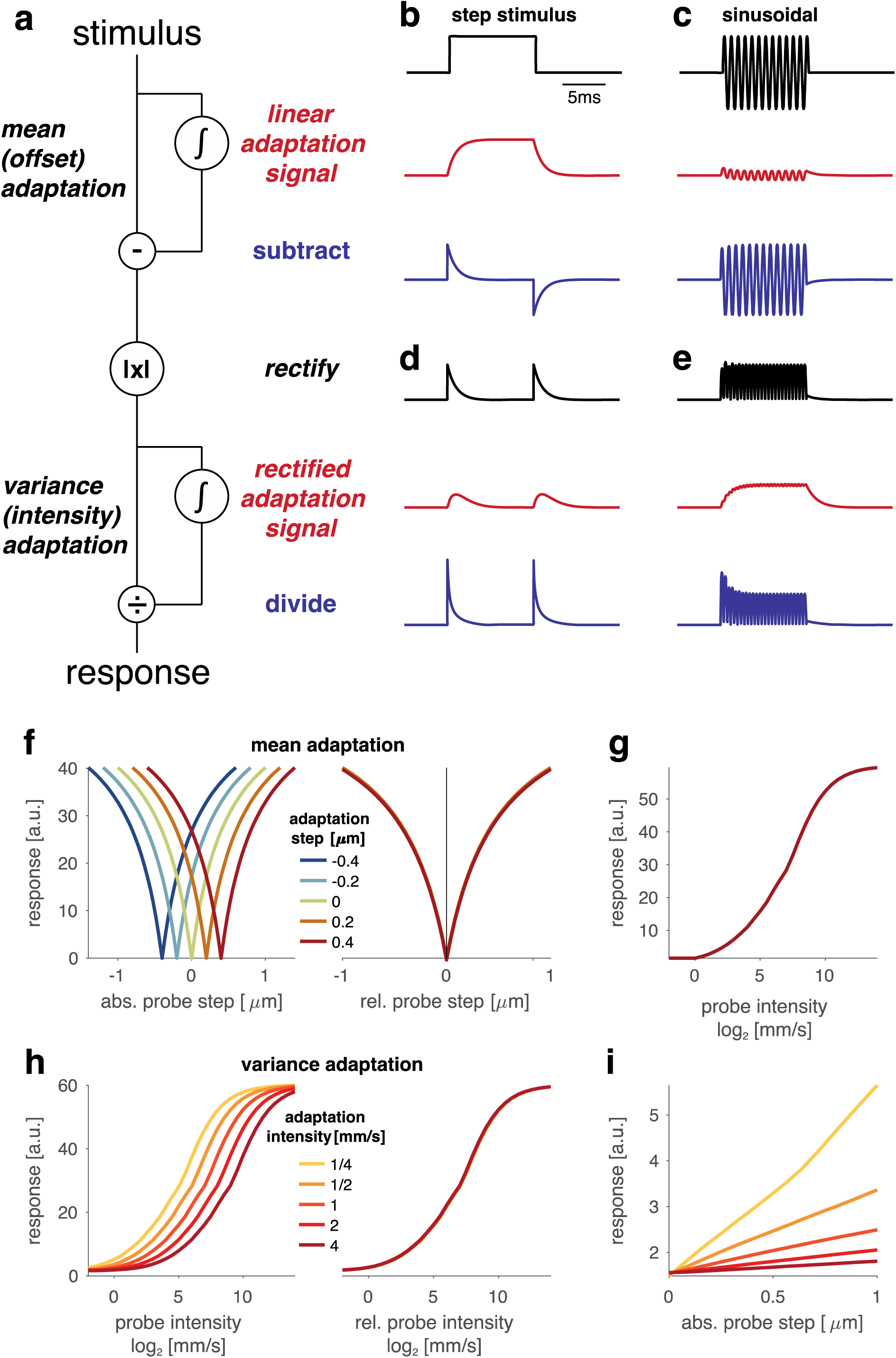
Computational model of adaptation in the JO. **a** A sequence of subtractive adaptation, rectification, and divisive adaptation reproduces our experimental data (see also Supplementary Fig. 11a). **b-e** Signals from different stages of the model in a for step (b, d, 10 ms) and sinusoidal (c, e, 400 Hz) displacement. Adaptation signals and adapted signals are colored red and blue, respectively. Shown are the stimulus (black in b and c), the linear adaptation signals (red in b and c), the output of the mean adaptation stage (blue in b and c), the rectified form of this output (black in d and e), the rectified adaptation signal (red in d and e), and the output of the variance adaptation stage corresponding to the overall output of the system (blue in d and e). **f** Step tuning curves of the model for different antennal baseline positions (adaptation steps, color coded, see legend) on an absolute scale (left panel) and relative to the baseline (right panel) (compare with data in Supplementary Fig. 10). The model qualitatively reproduces subtractive mean adaptation, which corrects for antennal baseline. **g** Intensity tuning curves of the model for different noise backgrounds (adaptation intensities, color coded, see legend) on an absolute intensity scale (left panel) and relative to the background intensity (right panel) (compare with data in Fig. 2b, c). Variance adaptation is divisive and compensates for sound intensity. **h** Intensity tuning of the model for different antennal baseline positions (color coded, see legend in d) corresponding to different levels of mean adaptation (compare with data in Fig. 5b). Intensity tuning is independent of mean adaptation. **i** Step responses of the model for different noise backgrounds (color coded, see legend in e) corresponding to different levels of variance adaptation (compare with data in Fig. 5e). Responses to steps are reduced with variance adaptation. The curves for different background noises or steps in d (right panel), e (right panel), and f are nearidentical and hence appear as single curves.

The output of the mean adaptation stage is then rectified (Fig. 6d, e), either through half-wave rectification (setting the negative parts of the signal to zero) or full-wave rectification (inverting the sign of negative signal parts) (Supplementary Fig. 12, left). Rectification transforms the standard deviation of the input (sound intensity) to the mean of the rectified output (Supplementary Fig. 12, right). This enables encoding of – and subsequently adaptation to – signal intensity by simple low pass filtering. Two features of the CAP responses suggest that JONs implement full-wave rectification: i) responses to step deflections are identical for positive and negative steps (Supplementary Fig. 10) and ii) responses to sinusoidals display frequency doubling (Fig. 3c, inset, compare Supplementary Fig. 12, bottom left). However, the specific form of rectification does not crucially affect adaptation in the model.

In the divisive adaptation stage (Fig. 6d, e), the adaptation signal now encodes stimulus intensity and divides the rectified signal to implement variance adaptation (Fig. 6h, compare Fig. 2b, c). Since the responses to steps have non-zero mean at the input of this stage (Fig. 6d), they also induce divisive adaptation, just as in our data, in which variance adaptation reduced step responses (Fig. 6i, compare Fig. 5e).

The model thus explains the unidirectional interaction between mean and variance adaptation in JONs: Mean adaptation does not affect sound responses (Fig. 6g), since the first, subtractive adaptation stage is basically blind to sound (Fig. 6c). By contrast, variance adaptation does affect step responses (Fig. 6i), since the adaptation signal in the second, divisive adaptation stage is non-zero for sounds *and* for steps (Fig. 6b, d). Overall, starting from our experimental findings on the interaction between mean and variance adaptation in the JO, our model proposes how efficient sensory adaptation to both mean and variance can be implemented.

## Discussion

Our analysis of adaptation in the *Drosophila* auditory system reveals that the receptor neurons subtract away changes in the physical stimulus mean, which corresponds to the baseline displacement of the sound receiver, and normalize for the variance of the stimulus, which corresponds to the sound intensity (Fig. 1-3). Both forms of adaptation first arise in JON subthreshold currents, before spiking (Fig. 4). Even though these two computations are implemented within a single neuron, they are partly independent: mean adaptation does not affect intensity tuning, but variance adaptation reduces responses to static receiver displacements (Fig. 5). A computational model suggests that this compartmentalization of mean and variance adaptation in JONs can be achieved by first subtracting the raw physical stimulus, then rectifying the mean-subtracted stimulus, and finally dividing by the rectified signal to correct for intensity (Fig. 6).

The two forms of adaptation studied here underlie an efficient representation of courtship song. As the female moves, her antenna is deflected by wind or gravity, while rapid changes in her position relative to the male singer induce fluctuations in song intensity at her ear. Mean adaptation has been shown to be subtractive and to render the auditory JONs largely insensitive to static or slow deflections^18,19^. Here, we show that variance adaptation is divisive and induces intensity-invariant responses after ~40ms. This contrasts with variance adaptation in the vertebrate ear, where the tuning curves of individual auditory nerve fibers do not shift sufficiently to correct for background intensity^47 48^.

Variance adaptation in the auditory receptor neurons, while fast enough to correct for the weakly amplitude modulated sine song, does not correct for the intensity of individual pulses within the pulse song (Fig. 3d, f, h). However, it still improves the representation of pulses, as it corrects for background sound levels (Fig. 2). In addition, variance adaptation underlies a high-pass filter for the inter-pulse interval (Fig. 3j), indicating that tuning for conspecific song features begins to arise at the level of the receptor neurons. Behavioral tuning for inter-pulse interval is a band-pass centered around values found in the conspecific song (35-40ms)^49-51^ and auditory neurons in the fly brain (AMMC-B neurons) already exhibit weak bandpass tuning for inter-pulse interval. Our results suggest that this tuning may in part be inherited from the JON inputs^52^. The incomplete adaptation to pulse trains containing *melanogaster*-specific inter-pulse intervals may be useful for encoding absolute pulse intensity, and this may facilitate orientation behaviors^53^. In addition, first-order auditory neurons in the fly brain (AMMC neurons) show adaptation across pulses in a train^9^, thereby further correcting for pulse intensity variation.

Information about antennal position in the sound-responsive type AB JONs is ambiguous since onset responses for step deflections depend on the intensity of the superimposed sound (Fig. 5e). However, type CE JONs do not adapt to antennal offsets and hence encode absolute antennal position^11,12,23^. Thus, just like in the mammalian somatosensory system, where different neurons adapt differentially to encode different aspects of touch^54^, distinct sensory channels in the *Drosophila* antenna differ in their adaptation properties to encode distinct aspects of antennal displacement.

Our data strongly suggest that adaptation is not a population-level phenomenon, but arises within individual type AB JONs. We recorded CAP signals from only type AB or B JONs (Fig. 1f), and recorded spiking and subthreshold activity from a subpopulation of type A JONs (Fig. 4d) – for both experiments we observed adaptation transients. We also recorded calcium signals from type A or B JONs (Supplementary Fig. 4) – unlike the CAP signal, calcium signals recorded via GCaMP6f are a measure of population firing rate not sensitive to spike timing; the existence of adaptation transients in the calcium signals further indicates that changes in JON sensitivity – not in population synchrony – underlie adaptation. Finally, simple models revealed that explanations for adaptation based on response inhomogeneity within the type AB JONs are unlikely to explain the observed adaptive CAP dynamics (Supplementary Fig. 3, 5, 6). These experiments and models do not rule out alternative explanations of adaptation in the CAP – however, they demonstrate that adaptation within individual type AB JON is the most parsimonious explanation of all of our data combined.

While the molecular bases for mean and variance adaptation in the JONs are still unknown, our data and model constrain hypotheses about their biophysical implementations. Mean and variance adaptation in JONs interact unidirectionally – mean adaptation does not affect sensitivity to stimulus variance but variance adaptation does affect responses to offsets (Fig. 5). Our model suggests that this can be implemented by a serial arrangement of two adaptation stages with an interleaved rectification step. The first adaptation stage subtracts the stimulus mean. After an interleaved rectification step, the second adaptation stage is divisive and corrects for intensity (Fig. 6). The partial compartmentalization of the two adaptation forms within JONs can be achieved by implementing mean adaptation at the mechanotransduction step. This is supported by mean adaptation affecting the mechanical sensitivity of the antenna and thus being visible in the antennal movement^18,21^. Mean adaptation is thought to occur via a stiffening of the gating springs that transform the antennal movement into opening and closing of mechanosensitive channels in JONs^18,54^. For mean adaptation to not affect sound sensitivity, the underlying adaptation signal must not be rectified, but must be a linear representation of the symmetrical antennal vibrations (Supplementary Fig. 11a).

Since variance adaptation is not visible in the antennal movement it likely arises downstream of the gating of mechanotransduction channels. We find that rectification compartmentalizes mean and variance adaptation in JONs, not a separation of timescales, since mean or variance adaptation are equally fast^18^ (Fig. 1e, Supplementary Fig. 10). Rather, the separation arises because i) the mean of the raw stimulus – prior to rectification – is independent of the variance and ii) rectification after mean subtraction transforms the stimulus variance to the mean of the rectified signal (Supplementary Fig. 12). Thus, simply dividing by the rectified, mean-subtracted stimulus implements variance adaptation^46^. The same principle holds for retinal adaptation to visual contrast^55^ and for intensity adaptation in hair cells^56^. Rectification may be implemented in JON dendrites via voltage-sensitive channels whose activation curve exhibits a hard or soft threshold^20,57,58^. Intriguingly, the frequency doubling in the CAP – a 300 Hz tone induces 600Hz CAP oscillations (Fig. 3c) – could be the result of full-wave rectification that transforms a sinusoidal’s negative lobes into positive ones (Supplementary Fig. 12). However, in the absence of single-neuron recordings of JONs, it is unclear whether this frequency doubling arises within single neurons or whether it is the outcome of recording from a population of neurons that respond at different phases of the sinusoidal^19^. Nevertheless, it is tempting to speculate that the frequency doubling in the CAP is a correlate of rectification for variance adaptation in single JONs.

Which molecules mediate variance adaptation? One possibility includes low threshold, voltage-sensitive potassium channels that open in response to subthreshold depolarizations with a delay and can thus dynamically reduce the amplitude of the generator currents (e.g.^59^). Adaptation channels can operate in a calcium-dependent manner, but we found that variance adaptation in *calmodulin* mutants appeared normal (Supplementary Fig. 9). Alternatively, the TRP channels *nompC*, *nanchung* and *inactive* have been shown to adjust the gain of mechanotransduction in the JO^37^. They could hence also play a role in variance adaptation which corresponds to a dynamical gain adjustment. However, we also found that *nompC* mutants display normal variance adaptation within their dynamic range (Fig. 4h, Supplementary Fig. 9). This is consistent with the observation that *NompC* mainly affects antennal gain and thus presumably acts upstream of variance adaptation^37^ (Fig. 4b). One major difficulty in discovering the molecules that mediate variance adaptation is that these molecules may also be involved in establishing the generator currents themselves (for example, *nanchung* and *inactive* mutants abolish the generator current^19^) – disentangling the two requires a way to measure adaptation and mechanotransduction independently.

How different forms of adaptation are organized in sensory receptor neurons fundamentally depends on the properties of the physical quantity they process. For the visual system, the mean and variance of a luminance pattern are both scaled by changes in illumination and hence can be corrected for in a single divisive step within photoreceptors^46,60^. Receptor neurons processing physical quantities that obey a similar scaling law between mean and variance – for instance those that correspond to a count or a rate – should organize adaptation in this way. The olfactory sensory neurons process the molecule count or odorant concentration, the mean and variance of which are similarly scaled by diffusion. Accordingly, olfactory receptor neurons have been shown to implement divisive –not subtractive – adaptation to concentration^61-64^. This also applies to adaptation in higher-order neurons, e.g. adaptation to visual or acoustic contrast^30,46,65^.

By contrast, the mean and variance of the physical quantity processed by auditory receptor neurons do not obey the same scaling law since they are typically affected by independent processes: the additive mean corresponds to a baseline displacement of the mechanical receiver, while the multiplicative variance corresponds to fluctuations in intensity of the actual sound stimulus, which is superimposed on the time-varying baseline. Adaptation in cochlear hair cells and now in JO neurons is organized accordingly, first subtracting the mean and then dividing by the variance^7,47^. All receptor neurons that process physical quantities with an additive mean and a multiplicative variance – e.g. quantities that can have both positive and negative values – should follow this pattern. While the existence of adaptation has been reported for virtually all sensory systems, the computations and interactions of various forms of adaptation are rarely studied. For instance, mechanosensory bristles in *Drosophila* implement subtractive adaptation to step deflections^66^ but to the best of our knowledge neither variance adaptation nor its interaction with mean adaptation have been studied. Likewise, primary sensory afferents of the whisker system adapt to different degrees of whisker deflection, but the computations underlying adaptation are unknown^67^. Strikingly, different afferents encode different aspects of whisker motion, like position or velocity^68^, and this diversity could be explained through the diversity of adaptation – e.g. strong mean adaptation supports encoding of velocity and prevents encoding of whisker position. Our combined experimental and modeling strategy for studying the conserved processes of mean and variance adaptation, and their interactions, can now be used to facilitate a genetic dissection of the molecules that underlie both forms of adaptation, in the *Drosophila* model system.

## Methods

### Flies

Group housed, virgin females were used for all recordings unless noted otherwise. Extracellular recordings were performed 2-5 days post eclosion. The wildtype strain used was *Canton S* (*CS*). Since homozygous *nompC* mutants are lethal, we used transheterozygous mutants for two different *nompC* null alleles (*nompC^1^/nompC^3^* and *nompC^3^/nompC^1^*, depending on whether the *nompC^1^* allele was inherited from the mother or the father) as in ^19^. Similarly, we generated transheterozygotes for the *cam* locus (*cam^n339^/cam^5^*) ^20^. We recorded CAPs from genetically identified subsets of JONs using flies mutant for *iav* (*iav^1^/iav^1^*) and we rescued the mutation by expressing wild type *Iav* under the control of JO1 (JON-ABCDE), JO2 (JON-B), or JO15 (JON-AB) ^19,69^. Calcium signals of JON-AB were recorded using GCaMP6f under the control of JO15-Gal4. Patch-clamp recordings from the giant fiber neuron (GFN) were performed in flies expressing eGFP in C17-GAL4. Since the generator currents were stronger in younger flies ^19^, we recorded at days 1-2 post eclosion. All flies were raised on a 12:12 dark-light cycle. *UAS-HM-iav* was provided by R. Wilson. All other stocks were provided by the Bloomington stock center.

Detailed genotypes used:

Fig. 1b-e, 2, 3, 5, Supplementary Figures 1b-c, 7, 8, 10, *Canton S*

Fig. 1f, Supplementary Figures 1d-e, *Canton S* (wild type) and *iav^1/^iav^1^* (iav^1^) and *iav^1^/iav^1^;UAS-HM-iav/+;JO1-Gal4/+* (JON-ABCDE) and *iav^1^/iav^1^;UAS-HM-iav/+;JO15-Gal4/+* (JON-AB) and *iav^1^, JO2-Gal4/ iav^1^, JO2-Gal4; UAS-HM-iav/+;+* (JON-B)

Fig. 4b-g, *Canton S* (antenna, CAP), *+; UASeGFP2x/C17-Gal4* (generator, spike)

Fig. 4h, Supplementary Figure 9, *Canton S* (wild type) and *+;nompC^1^,cn,bw/cyo* (nompC^1^/cyo), *+;nompC^3^,cn,bw/cyo* (nompC^3^/cyo) and *+;nompC^1^,cn,bw/nompC^3^,cn,bw* (nompC^1^/nompC^3^ (maternal/paternal) and nompC^3^/nompC^1^ (maternal/paternal) and cam^n339^/cam^5^ (Supplementary Fig. 9 only)

Supplementary Fig. 1a *+;JO15-Gal4/UASeGFP2x* (patch clamp)

Supplementary Fig. 4, *+;UAS20x-GCaMP-6f-UAS-Tom/cyo; JO15-Gal4/Tm6,tb* (Calcium imaging)

### Stimulus design and presentation

Stimuli were generated at a sampling frequency of 10 kHz. Band-limited Gaussian noise (from now on termed "noise”) was produced from a sequence of normally distributed random values by band-pass filtering using a linear-phase FIR filter with a pass band between 80 Hz and 1000 Hz. For probing adaptation, we switched the intensity of the noise every 100 ms (or 1000 ms for Ca^++^ imaging) in a sequence that contained all transitions between ¼, ½, 1 and 2 mm s^-1^ (Fig. 1b). The effect of adaptation on intensity tuning was assessed using a noise stimulus at intensities ¼, ½, 1, 2, 4 mm s^-1^ (adaptation background) whose intensity was switched every 120 ms for 20 ms to a probe intensity of 1/16, 1/8, ¼, ½, 1, 2, 4, 8 mm s^-1^ (e.g. Fig. 1F). To minimize artifacts from abrupt changes in sound intensity, each intensity switch had a duration of 1 ms during which the intensity was linearly interpolated to the new value.

Artificial pulse song (Fig. 3d) was generated as a train of Gabor wavelets: sin(x*2πf_c_+φ) exp(-(x/σ)^2^ with carrier frequency fc of 250 Hz, phase φ of 0π, standard deviation σ of 4.6 ms and an inter-pulse interval of 36 ms. These parameters were chosen to mimic the shape of song pulses produced by *Drosophila melanogaster* males during natural courtship^33^. For Fig. 3j, the inter-pulse interval was varied between 8 and 72 ms.

#### Sound

The sound delivery system consisted of i) analog output of a DAQ card (PCI-5251, National Instruments), ii) a 2-channel amplifier (Crown D-75A), iii) a headphone speaker (KOSS, 16 Ohm impedance; sensitivity, 112 dB SPL/1 mW), and iv) a coupling tube (12 cm, diameter 1 mm).

The stimulus presentation setup was calibrated as in ^9^. Briefly, the amplitude of pure tones of all frequencies used (100-1000 Hz) was set using a frequency-specific attenuation value measured with a calibrated pressure gradient microphone (NR23159, Knowles Electronics Inc., Itasca, IL, USA). To ensure that the temporal pattern of the noise stimuli was reproduced faithfully, we corrected the presented noise patterns by the inverse of the system’s transfer function, measured using a pressure microphone (4190-L-001, Brüel & Kjaer). During experiments, the sound tube was positioned at a distance of 2 mm and an angle or 90° to the right (extracellular recordings and laser Doppler vibrometry) or left (Calcium imaging) arista coming from behind the fly.

#### Piezoelectric actuation

Piezoelectric actuation was performed using a single channel piezo driver (MDT694, Thorlabs) and actuator (A186). The piezo actuation was calibrated using laser Doppler vibrometry (see below) by measuring displacement and velocity of the piezo tip for all driving voltages used in the experiments (steps and 300 Hz sinusoidals). Step stimuli for piezo-electric actuation were low-pass filtered using a 1ms Gaussian window to stay within the operating limits of the piezo driver.

### Electrophysiology

Extracellular recordings were performed using glass electrodes (1.5OD/2.12ID, WPI) pulled with a micropipette puller (Model P-100, Sutter Instruments). The fly’s wings and legs were removed under cold anesthesia and the abdomen was subsequently fixed using low-temperature melting wax. The head was fixed by extending and waxing the proboscis’ tip. The preparation was further stabilized by applying wax or small drops of UV-curable glue to the neck and the proboscis. The recording electrode was placed in the joint between the second and third antennal segment and the reference electrode was placed in the eye. Both electrodes were filled with external saline ^70^.

The recorded signal was amplified and band-pass filtered between 5 and 5000 Hz to reduce high frequency noise and slow baseline fluctuations induced by spontaneous movement of the antenna (Model 440 Instrumentation Amplifier, Brownlee Precision). We ensured that the band-pass filter did not distort the recorded signal, e.g. introduce artefactual response transients. We subsequently digitized at 10 kHz with the same DAQ card used for stimulus presentation (PCI-5251, National Instruments). Patch clamp recordings from the giant fiber neuron were performed as described in ^9,70^. After obtaining access to the membrane voltage (input resistance between 70 and 400 MΩ), we recorded in voltage clamp mode the spike-induced current, or – after bath application of TTX (Tocris, UK) to 6 μM final concentration – the generator current.

### Calcium imaging

Calcium signals were recorded from flies expressing GCaMP6f ^28^ under the control of JO15-Gal4, which expresses in the majority of JON-AB ^69^. Flies were anesthetized on ice and gently waxed into the hole of a perfusion chamber with the proboscis facing upwards such that both antennae were free to move on the underside of the chamber. The antennal nerves and the AMMC were exposed by removing the proboscis and surrounding tissue. Sound stimuli were presented using sound delivery tubes as described above.

Two-photon imaging was performed using a custom microscope controlled by Scanimage software (Vidrio Technologies). The calcium indicator was excited at 940 nm using a TiSa Laser (Coherent). GCaMP emission were detected using a resonant scanner at a frame rate of ~60 Hz. ROIs were drawn manually around the projections of type A and type B JONs in AMMC ^69^ (Supplementary Fig. 4a). Calcium responses from both JON populations were similar (Supplementary Fig. 4c) and we pooled data from both populations for each fly for all analyses (Supplementary Figures 4d,e). ΔF\F values were obtained by subtracting the average fluorescence in the second immediately prior to sound stimulus onset.

### Laser-Doppler vibrometry

Arista movement was measured using laser-doppler vibrometry (Polytec OFV534 laser unit, OFV-5000 vibrometer controller, Physik Instrumente, low-pass 5 kHz).

### Data analysis

#### Pre-processing

CAP/LVM instantaneous amplitude was estimated as the magnitude of the Hilbert transform. This method is preferable over "root mean square” like algorithms ^71^ since it allows estimating the amplitude without imposing a time scale through the duration of the smoothing time window.

#### Adaptation time scale and strength

The adaptation time scale was estimated by fitting an exponential function *r(t)=r_0_ + r_max_exp(-t/τ)* to the falling/rising phases of positive/negative transients (Fig. 1c, 3f). Adaptation strength (Fig. 3g) was estimated based on the slopes s_on_ and s_ss_ of the onset and steady state intensity tuning curves (e.g. Fig. 1d), e.g. *s_on_=<Δr_on_(*x*)Δx>_x_* where <.>_x_ denotes the average over intensities. *1-s_on_/s_ss_* is a measure of adaptation strength that is independent of response magnitude: It’s 0 for no adaptation (*s_on_=s_ss_*), approaches 1 for complete adaptation (*s_ss_≪s_on_*), and can exceed either of these values in the presence of noise.

#### Sigmoidal fits of the tuning curves

Onset and steady responses for the intensity-step stimuli (Figures 1b-d) and for the background-adaptation stimulus (Fig. 1f, onset only) were obtained by averaging the CAP amplitude over the first 8ms and last 10ms of a step/probe, respectively. The tuning curve shapes did not depend crucially on the specific method chosen – results were similar when using different averaging time windows or when taking the CAP amplitude extrema instead.

Sigmoidal fits to the resulting curves were obtained using the least squares method. We extracted the slope *α* and offset or shift *β* of the tuning curves from these fits: *r(x)= r_0_ +r_max_/(1+e xp(-α(x-β))*, where *x* is the sound intensity on a logarithmic scale and *r(x)* is the CAP amplitude. Fits were excellent throughout (r^2^>0.9).

#### Fisher information

Fisher information (*I_fisher_*) is a local measure of stimulus (intensity) discriminability that can be estimated from the tuning curves. *I_fisher_ increases* with the magnitude of the local slope of tuning curve (Δr\Δχ)^2^ – steeper slopes produce larger response changes for a given stimulus change and hence facilitate stimulus discrimination. *I_fisher_ decreases* with the local variance of tuning curve σ^2^(x) since variability adds uncertainty to stimulus induced response differences^29^. Tuning curve slope as a function of intensity was estimated by taking the derivative of sigmoidal fits to the experimental tuning curves (see above). Since in our data, the response variance is proportional to mean (r^2^=0.98), Fisher information is proportional to *I_fisher_(x) ∝(Δr(x)/Δx)^2^/r(x)* ^72^.

#### Statistical methods

In all figures, error bars depict mean ± std over animals. For the statistical tests in Figures 3f and g we chose a nonparametric test (sign test for paired, rank sum for unpaired data) since data points were not normally distributed (based on a Jarque-Bera test for normality). Z-scores for the tests are 2.27, 0.65 in Fig. 3g and -2.45, -2.04, -2.65 in Fig. 3f (left to right). P-values in Figures 4c and f derive from the significance of the r^2^ value of a linear fit to the values for each fly (per fly p-values and t-statistics (in parentheses) for Fig. 4c: 0.693 (-0.44), 0.139 (1.99), 0.769 (-0.32), 0.158 (-1.86), Fig. 4f: 0.0091 (10.41), 0.0096 (10.16), 0.0003 (57.59), 0.0066 (12.29), 0.0065 (12.30), 0.0050 (14.04), 0.0085 (10.80)). 95% confidence intervals (CIs) in Figures 1f and 2g, h were estimated as two times the standard error of the mean over trials or animals, respectively.

### Proof-of-principle models

To explore different explanations for the adaptive dynamics in the CAP, we employed regular and adaptive leaky integrate and fire (LIF) model neurons ^73^.

The subthreshold dynamics of the membrane voltage (V_m_) in a LIF is given by: τ_V_d/dt V_m_(t) = I_in_(t) – I_leak_(t) – I_adapt_(t), with input current I_in_ = σ_stim_I_stim_(t) + σ_noise_I_noise_(t) and additive noise I_noise_. The leak current I_leak_ with membrane resistance R_m_ follows I_leak_(t) = V_m_(t) – V_rest_. The dynamics of the adaptation current I_adapt_ evolve according to τ_adapt_ d/dt I_adapt_(t) = I_adapt_(t). A spike is elicited when V_m_ reaches the threshold V_thres_ after which it is reset to V_res_. I_adapt_ increments by ΔA upon each spike and is 0 for a regular LIF. For all simulations τ_v_ = 4 ms, V_thres_ = 1 mV, and Vrest = -65 mV, except for Supplementary Fig. 3c, for which the tv was chosen randomly from the interval 2 and 8 ms. The time constant of the adaptation current τ_adapt_ is 25 ms for Supplementary Figures 2d, 3d and 100 ms for Supplementary Fig. 5e-f. The latter time constant is much longer than observed in the CAP. This value was chosen to be able to observe an effect of adaptation for this stimulation paradigm by enabling adaptation to carry across subsequent steps in intensity. For the regular LIF models (Supplementary Figures 2b-c, 3b-c, 5c-d), the adaptation strength ΔA is 0 mV. For adaptive LIF models, ΔA is 1 mV for Supplementary Figures 2d and 5e-f, and 1. 5 mV for Supplementary Fig. 3d.

Stimuli were defined in terms of sound intensity (mm s^-1^) and then scaled to an input current Istim to obtain firing rates that were comparable across conditions (regular vs. adaptive LIF models) and physiologically plausible. Noise stimuli were low pass filtered white noise (cutoff 1000 Hz) with step changes in intensity (standard deviation of the white noise) as describe in the figure legends. For Supplementary Fig. 3, the noise had a mean of 2 mm s^-1^ to induce constant depolarization and to promote spike time jitter to accumulate over time. For Supplementary Fig. 2b, the noise stimulus was identical for all neurons and independent white noise was added for each neuron to introduce spike-time jitter. For all other simulations, the stimulus was a different realization of the noise since we were interested in observing changes in spike timing and firing rate associated with the steps in intensity, not with the temporal pattern of the white noise. For the range fractionation model (Supplementary Fig. 5), the low-intensity population (blue) was fed the noise stimulus only while the noise was at low intensity (1 mm s^-1^) and the high-intensity population (orange) only received input during the high-intensity period (2 mm s^-1^). LIF models were numerically simulated using Euler’s method with a step size dt=1/20 ms.

### Model of JON adaptation

Subtractive and divisive adaptation were modeled as an incoherent feedforward loop ^45^ which consists of two steps: First, leaky integration low-pass filters the signal to an adaptation signal *x_ada_* = ∫ *x_in_ exp*(*-t/τ*)*dt*. The integration time constant *τ* controls how fast the adaptation signal is updated and hence determines the effective speed of adaptation. Second, the adaptation signal then either subtracts or divides the input: *x_out_ = x_in_ - x_ada_* or *x_out_ = x_in_/(1/σ_div_ + x_ada_). 1/σ_div_* is added to the adaptation signal in the divisive case to avoid division by zero in the absence of stimulation and to control completeness of adaptation.

Generally, *x_ada_* ≈*0* for stimulus symmetrical fluctuations much faster than *τ*. Hence there is no adaptation for rapidly fluctuating and symmetrical stimulus components, i.e. sound-induced vibrations, and adaptation acts as a high-pass filter. Note that this model can in principle also account for power law dynamics in adaptation ^74,75^, by replacing the exponential integration kernel with a power law kernel ^76^. This does not affect the qualitative model behavior we were interested in here. Rectification in the model was implemented as either full-wave rectification *y*=|x| or half-wave rectification *y*=*Θ(x)*, where *Θ(x)=0* for *x<0* and *Θ(x)=x* for *x≥0* (Supplementary Fig. 12).

We systematically explored the adaptation properties of all serial and parallel arrangements of the three elementary computations: subtractive adaptation (S), divisive adaptation (D), rectification (R). To that end we generated computational networks using the following rules: For serial networks, the root node could have either R or Ø (“Ø” stands for an empty computational node) followed by an adaptation node containing S or D or Ø, followed by R or Ø, followed by another adaptation node containing S or D or Ø, followed by R or Ø. More compactly: R/Ø→S/D/Ø→R/Ø→S/D/Ø→R/Ø (“/” stands for “or”). These rules yielded 22 unique serial networks. Parallel networks contained two branches (rules for each branch: →R/Ø→S/D/Ø→R/Ø→) and an input and an output node (rules for each: →R/Ø→). This resulted in 100 parallel networks. The parameters for testing all 122 computational network motifs were: τ_sub_=30 ms, τ_div_=50 ms, σ_div_=10^-4^.

We tested for robustness of the qualitative model behavior to changes in model parameters by running the selected network motif (Fig. 6a, subtractive adaptation→rectification→divisive adaptation) for combinations of τ_sub_ and τ_div_ on a grid (10≤τ≤ 100, 20×20 values, linearly spaced, σ_div_=10^-4^, Supplementary Fig. 11b) and different values of σ_div_ (10^-8^≤σ_div_≤ 10^0^, 20 values, logarithmic spacing, τ_sub_=30 ms, τ_div_=50 ms, Supplementary Fig. 11c). The resulting tuning curves (compare Figures 6f-i) were fitted to sigmoidal functions (see above) to extract the change in slope and position with adaptation (top and bottom panels, respectively, in Supplementary Figures 11b-c). We then used the coefficient of variation (CV) across adaptation stimuli of the tuning curves’ shift and slope to quantify adaptation for each parameter combination and for the four adaptation paradigms employed in the paper. A model qualitatively matching our data should have the following properties: Due to mean and variance adaptation, steps and intensity are encoded relative to an adaptation step/intensity. For mean adaptation, tuning curve offset and slope are identical for different offsets if plotted in a x-scale relative to the adaptation step (Supplementary Fig. 10c). Hence, the CV across adaptation parameters should be close to zero. The same holds for intensity adaptation – tuning curves are virtually identical on a log intensity scale relative to the adaptation intensity (Fig. 2c). Accordingly, CV of tuning curve offset and slope should be small. For the impact of intensity adaptation on step responses, we assessed tuning curve parameters on an absolute probe scale – in our data, only the slope, but not the offset of the curves changes with different adaptation intensities (Fig. 5e) and hence the CV of offsets should be small and that of slope should be high. For the impact of mean adaptation in intensity tuning, we observed no change in intensity tuning for different adaptation step sizes (Fig. 5b) – accordingly, CV of tuning curve offset and slope should be near zero for a model that matches our data. The chosen network motif (Fig. 6a) matches these expectations (Figures 6f-i). Moreover, model behavior is robust to changes in all three parameters (see Supplementary Figures 11 b,c), except for small σ_div_, simply because this parameter controls adaptation strength and when adaptation is weak, tuning curves don’t change much.

### Data and code availability

All data and source code are available from the authors upon request.

## Acknowledgements

We thank Eero Simoncelli, Jonathan Pillow, Martin Göpfert, and Pip Coen for comments on the manuscript. We also thank Xiao-Juan Guan for technical assistance, and Brendan Lehnert for advice on GFN recordings. JC was funded by the DAAD (German Academic Exchange Foundation) and the Sloan-Swartz Foundation. MM is an HHMI Faculty Scholar and was also funded by a National Science Foundation CAREER award, NIH New Innovator Award, NSF BRAIN Initiative EAGER award, the McKnight Foundation, and the Klingenstein-Simons Foundation.

## Author contributions

J.C. and M.M. designed the experiments. J.C. and N.O.-E. performed the experiments. J.C. analyzed the data and designed the models. J.C. and M.M. wrote the paper.

## Conflict of interest

The authors declare no conflict of interest

---

**Supplementary Figure 1:**
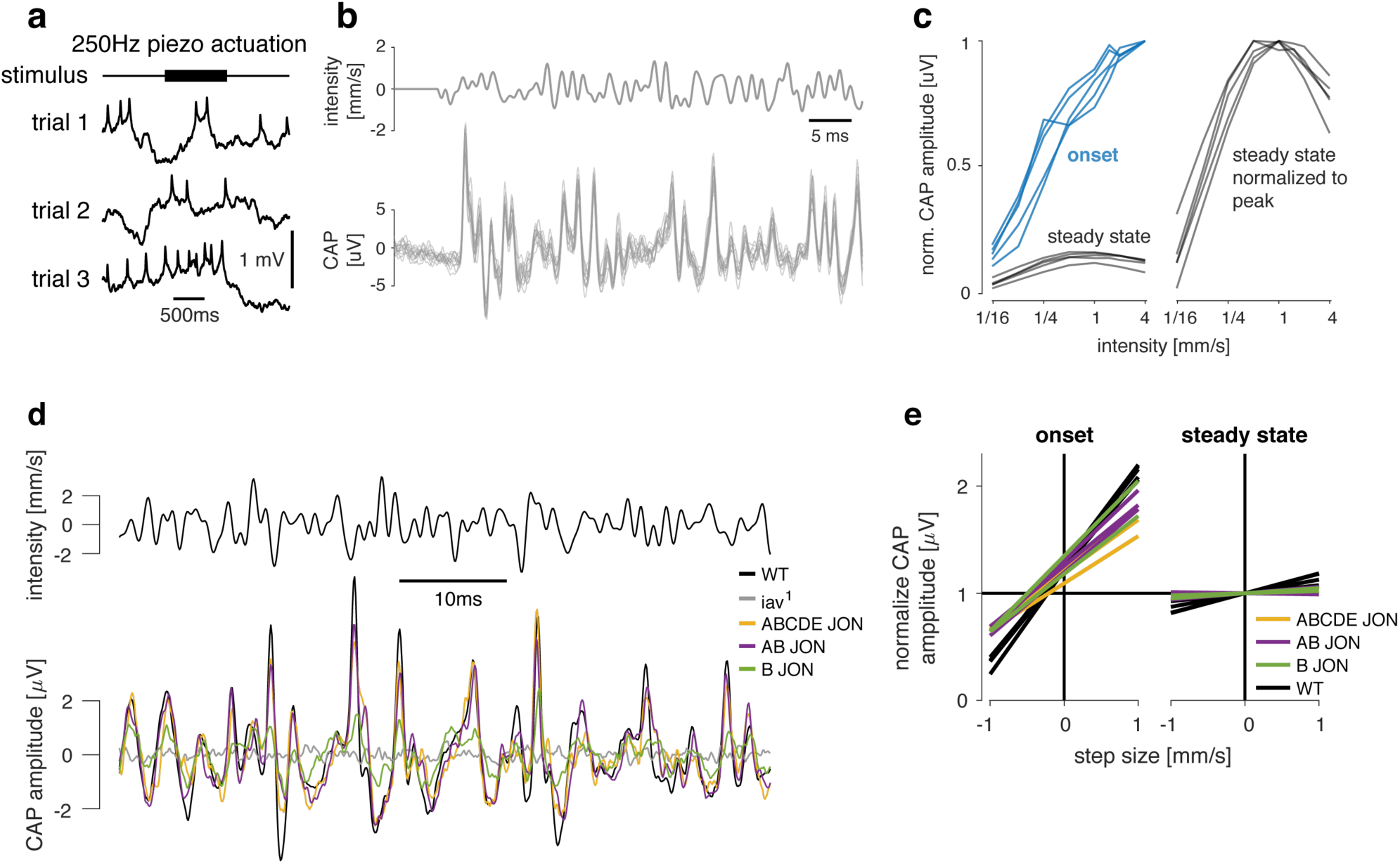
CAP recordings reflect firing of type A and B JON. **a** Patch-clamping interferes with mechanotransduction. Shown are 3 trials of a patch-clamp recording from the cell body of a type AB JON, during which the submerged arista was vibrated via piezoelectric actuation but no consistent, stimulus-evoked change in the membrane voltage was observed (stimulus = 250Hz, upper trace). **b** Single-trial responses (bottom) to a noise stimulus (top) are reliable from trial-to-trial. **c** Onset (blue, left) and steady-state (black, left) CAP amplitudes for noise responses for individual flies (N=5, averaged over 20 trials each). Curves for each fly were normalized to the maximal onset response. Right panel shows the steady tuning curves individually normalized to their respective maxima. **d** CAP responses (bottom) to a white noise stimulus (top) for wild type flies (CS Tully, black), *iav^1^* mutants (grey) and rescues in all (ABCDE) JONs (yellow), type AB JONs (purple), or type B JONs (green). The *iav^1^* mutants do not produce CAP responses. CAPs for rescues of all and AB JONs are similar to the wild type. CAPs of type B JONs rescues are weaker and noisier than the other rescues, suggesting that both A and B JON subpopulations contribute to the CAP for white noise stimuli. **e** Average onset (left) and steady-state (right) responses for all genotypes in Fig. 1f (N=2-5 flies per genotype, 20 trials each). Reponses are normalized for each fly such that the average steady-state response across step sizes is 1.0. Onset responses depend strongly on intensity and steady-state responses are near intensity invariant, independent of genotype (see legend).

**Supplementary Figure 2:**
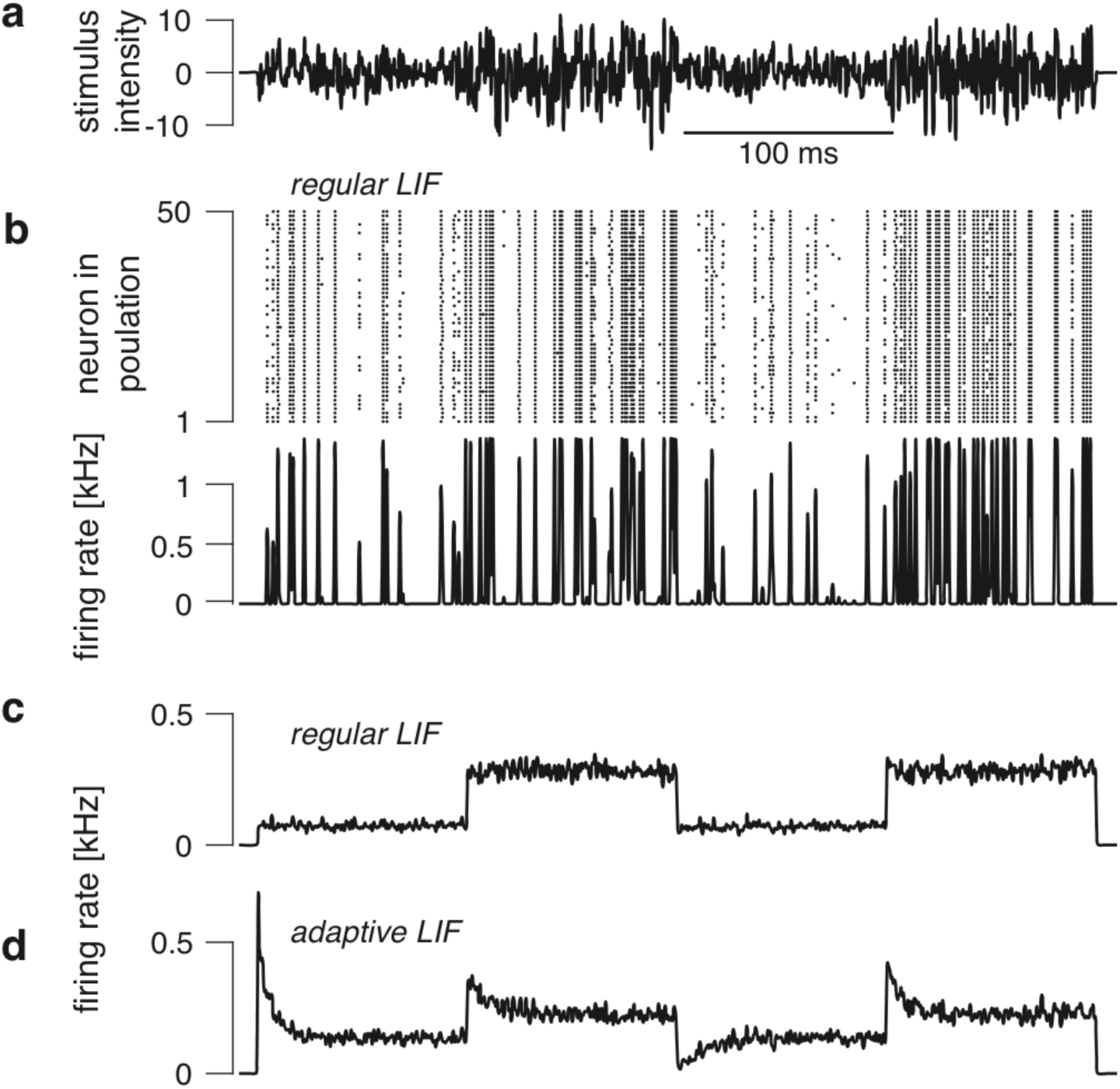
De-synchronization or response heterogeneity in JONs does not explain adaptation dynamics. We modelled a population of integrate and fire neurons with membrane noise to explore the effects of desynchronization (b-c) and of an adaptation current (d) on firing rate dynamics. For all panels, the neurons in each population had identical parameters and stimulus inputs and only differed in their membrane noise. See Methods for more details on the simulations. **a** Noise stimuli used in our study prevent strong de-synchronization in a population of leaky integrate and fire (LIF) neurons. Noise stimulus with zero mean and step-like changes in intensity (switches between 2 and 4mm s^-1^) as used in the CAP recordings and Ca imaging experiments. **b** Responses to the stimulus in a of a population of 500 LIF neurons, each with independent intrinsic noise. Spike raster plot (top, only a subset of 50 neurons shown for clarity) and population firing rate (bottom) do not exhibit de-synchronization at stimulus onset and no transients in synchrony or firing rate for changes in stimulus intensity following stimulus onset. Note that synchrony does change with stimulus intensity through an increase in the signal-to-noise ratio: louder stimuli produce stronger inputs relative to intrinsic noise sources and hence weaker desynchronization. However, this effect is not transient and hence does not explain adaptation. **c** Population average firing rate of LIF neurons for 500 independent realizations of the noise stimulus in a. There are no transients in firing rate. **d** Same as c but using LIF neurons with an adaptation current. Negative and positive transients arise upon changes in noise intensity just as in our CAP recordings.

**Supplementary Figure 3:**
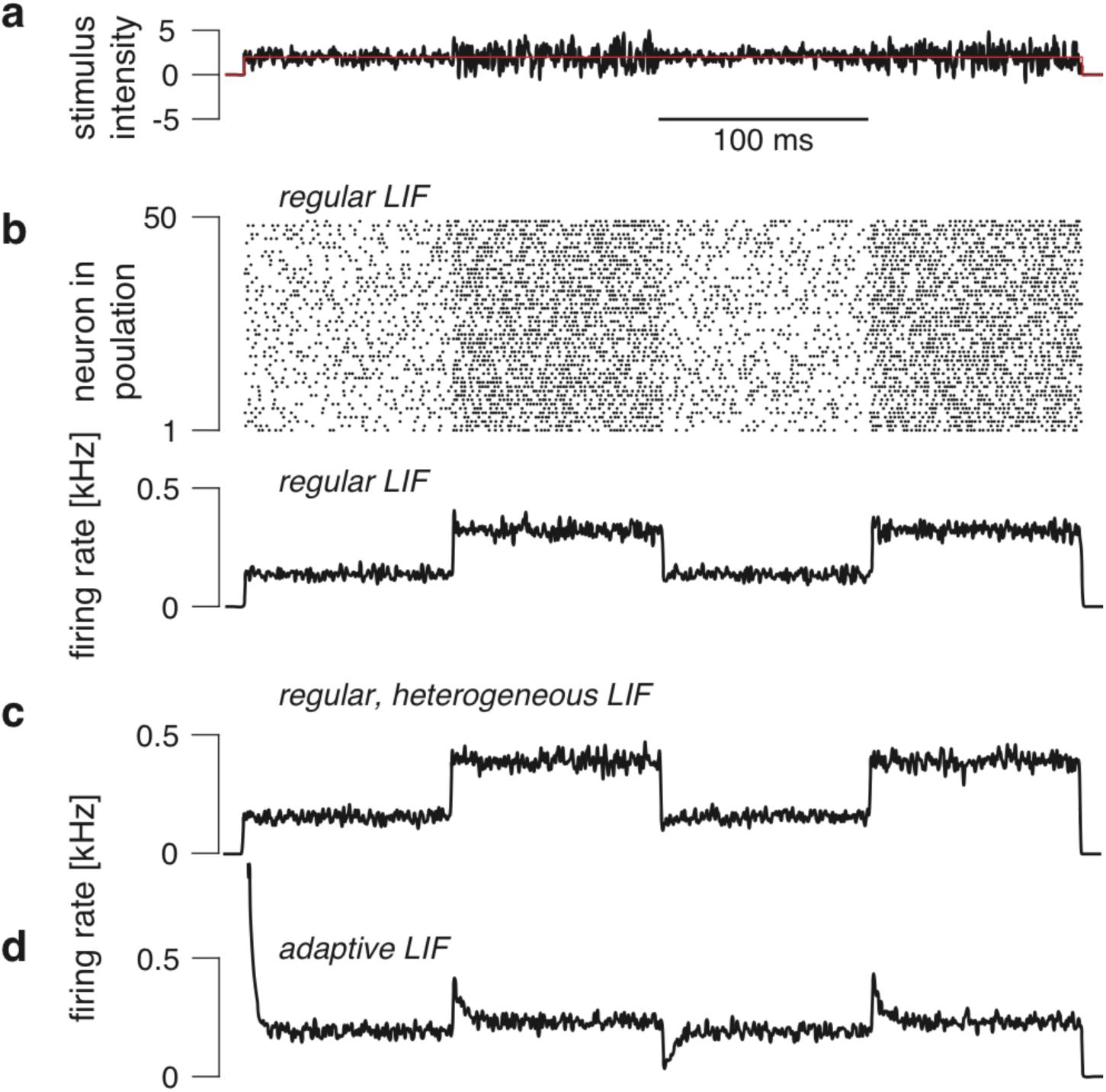
A stimulus regime that promotes de-synchronization or response heterogeneity in JONs does not explain adaptation dynamics. We modelled a population of integrate and fire neurons with membrane noise to explore the effects of desynchronization (b), response heterogeneity (c), and of an adaptation current (d) on firing rate dynamics. In contrast to the simulations shown in Supplementary Figure 2, we used noise with a constant offset as a stimulus to promote desynchronizations. For all panels except for panel g, the neurons in each population had identical parameters and stimulus inputs and only differed in their membrane noise. For panel g, we explicitly tested the effect of response heterogeneity by choosing the membrane time constant for each neuron in the population at random. See Methods for more details on the simulations. **a** Noise stimulus with step-like changes in intensity (switches between 0.5 and 1 mm s^-1^) with an added 2mm s^-1^ offset that induces a constant depolarization of the model neurons and promotes de-synchronization. **b** Responses of a population of LIF neurons for 500 independent realizations of the noise stimulus. Spike raster plot (top, only a subset of 50 neurons shown for clarity) and population firing rate (bottom) lack transients in firing. **c** Same as b but for a population of regular LIF neurons with heterogeneous adaptation and integration properties (membrane time constants in each neuron was scaled by a random factor between *½* and 2). The heterogeneity does not produce the strong transient changes in firing rate upon step changes in intensity see on the data. **d** Same as b but for LIF neurons with an adaptation current. Adaptation induces transient changes in firing rate – but not in synchrony – upon step changes in intensity. Overall this suggests that neither desynchronization nor population heterogeneity are not sufficient to explain the adaptation dynamics in the CAP and that adaptation is necessary to do so.

**Supplementary Figure 4:**
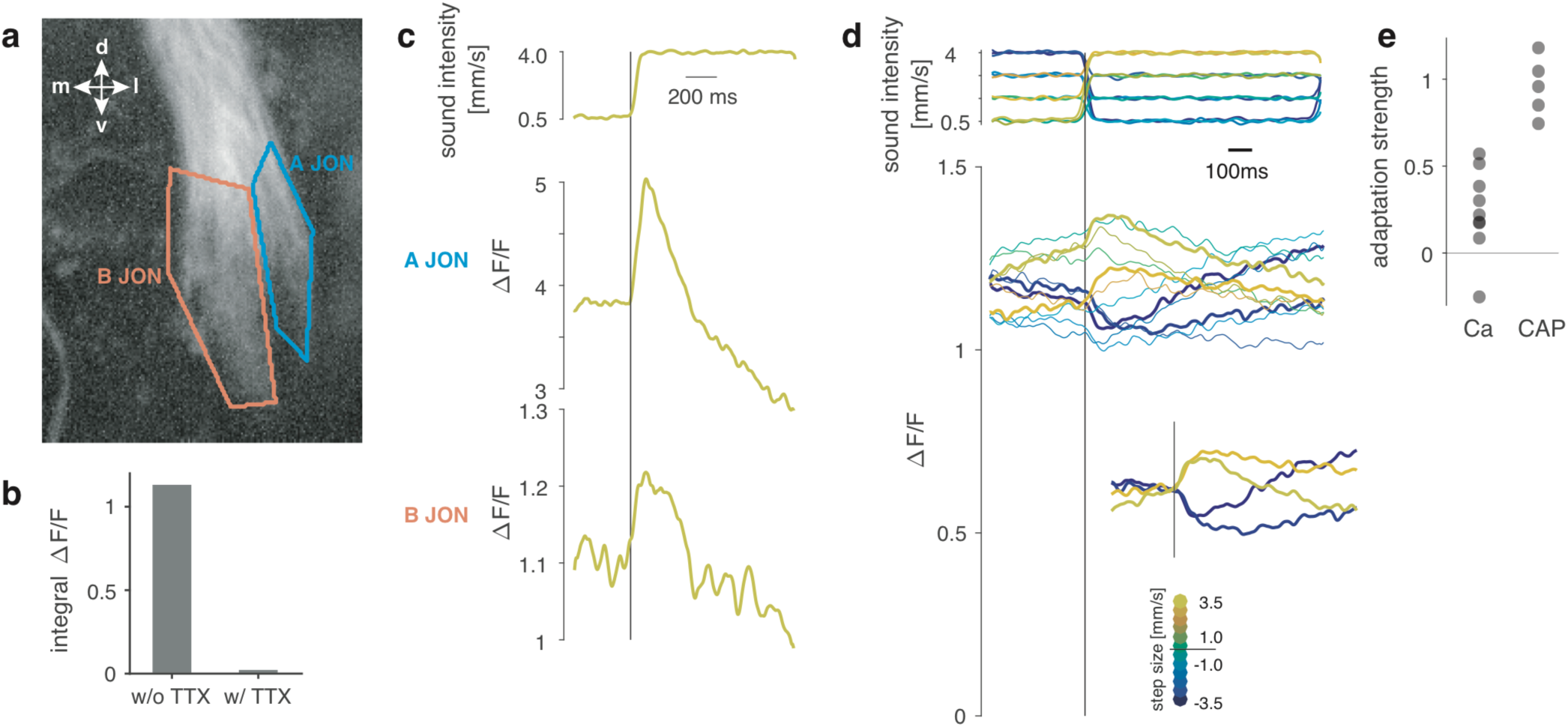
Calcium signals from type A and B JON display adaptation. **a** Type A and B JONs can be distinguished anatomically because they branch separately into the AMMC. Figure shows two-photon baseline fluorescence of JON expressing GCaMP6f under the control of JO15-Gal4. **b** Calcium responses (ΔF/F) to 200 Hz sinusoids with and without bath application of TTX (~6 μM). TTX has been shown previously to abolish spiking in JONs ^1^. c Representative calcium responses (ΔF/F) to the largest positive step in noise intensity measured in the projections of type A JONs (middle) or type B JONs (bottom) shown in panel a (40 trials per step). The stimulus is similar to that used for the CAP recordings (Fig. 1 b) but with an interval between steps of 1000ms to account for the slow dynamics of calcium signals. Both Type A and B JON projections produce transient changes in fluorescence upon step changes in noise intensity, indicating that both subpopulations adapt to sound intensity. These calcium transients are weaker and slower than those seen in the CAP (see Fig. 1), likely due to filtering by the calcium sensor. **d** Steps in noise intensity (top) used to probe adaptation in calcium signals. Responses are pooled across type A and B JONs in one fly (bottom). Step size is color-coded (see legend). The calcium signals (bottom) display weak but clear positive and negative transients locked to the time of the intensity step (traces from one fly, averaged over 20 trials per step). This is most evident for the responses to the two most positive and the two most negative steps in noise intensity (thick lines). To highlight the transients, the inset shows responses to the greatest steps in intensity, with baseline subtraction. Changes in fluorescence only weakly depend on absolute noise intensity, consistent with our observation of intensity invariance in the CAP. **e** Strength of adaptation was obtained by comparing onset responses to intensity steps to the responses at steady-state (see Methods for details, N=8 flies). Adaptation values of 1.0 indicate complete adaptation. Values for the CAP are reproduced from Fig. 4g.

**Supplementary Figure 5:**
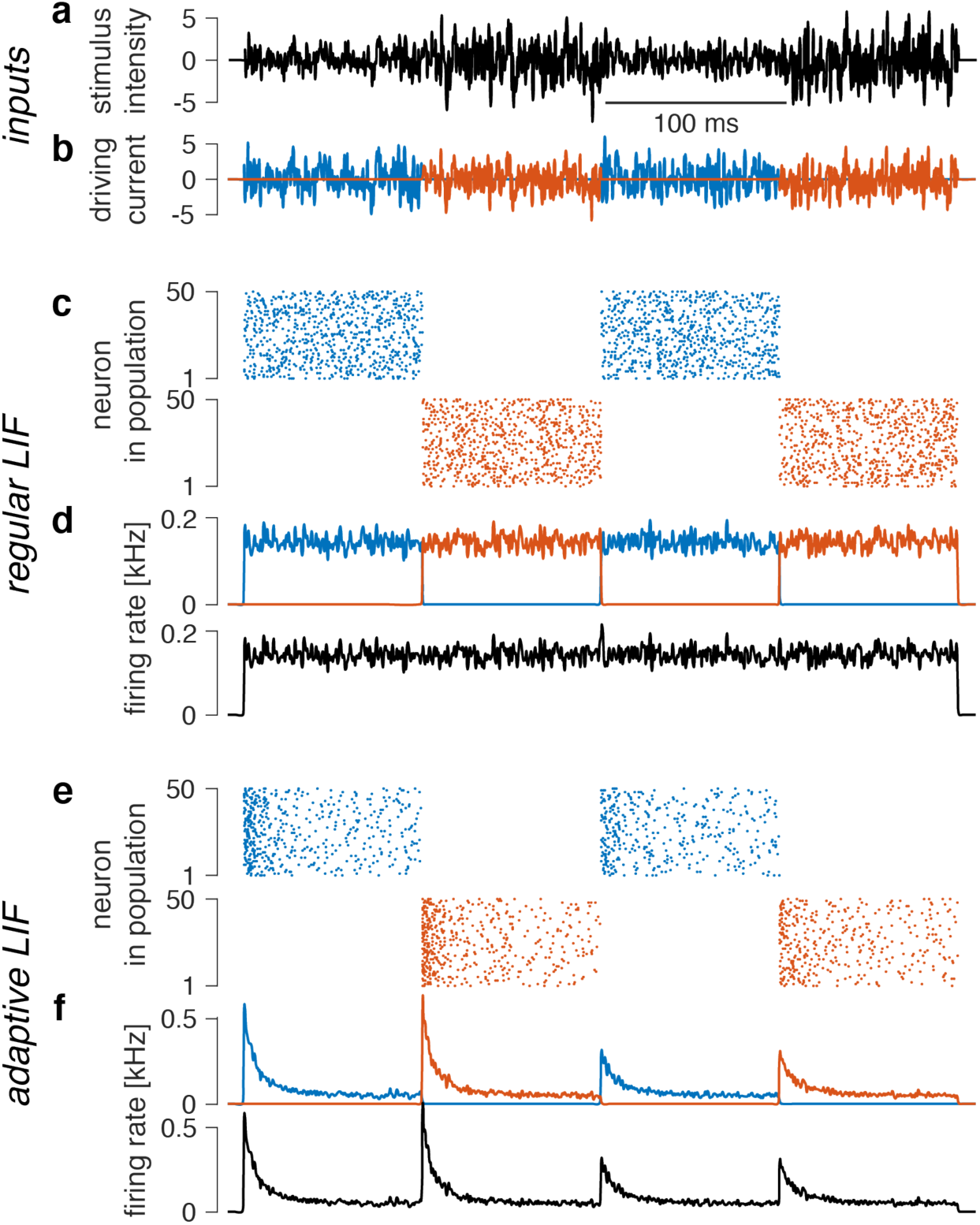
Intensity range fractionation does not explain adaptation dynamics. Heterogeneity in intensity tuning among individual JO neurons can produce intensity invariance at the population level through range fractionation – individual JONs tiling the relevant intensity range **a** Noise stimulus with steps in intensity (1 and 2 mm s^-1^, top). **b** Two populations of neurons that respond exclusively at noise intensities of either 1 (blue) or 2 (orange) mm s^-1^. **c, d** Spike raster plots (c) and population firing rates (d) of the low-intensity (blue) and high-intensity (orange) population of leaky integrate and fire (LIF) neurons. This model only produces transient increases in firing rate upon increases as well as decreases in intensity. By contrast, JONs produce positive transients for intensity increases, and negative transients for intensity decreases (Fig. 1c). **e, f** Same as c, d but with adaptive LIF model neurons. Adaptation attenuates the positive transients for the two last steps in noise intensity if the adaptation time constant exceeds the step duration. However, adaptation fails to reproduce the pattern of positive and negative transients in the population firing rate observed in the CAP. Intensity range fractionation – or more generally heterogeneity in intensity tuning among JONs - is thus unlikely to underlie the adaptive dynamics of the CAP. See Methods for details on the parameters of the simulations.

**Supplementary Figure 6:**
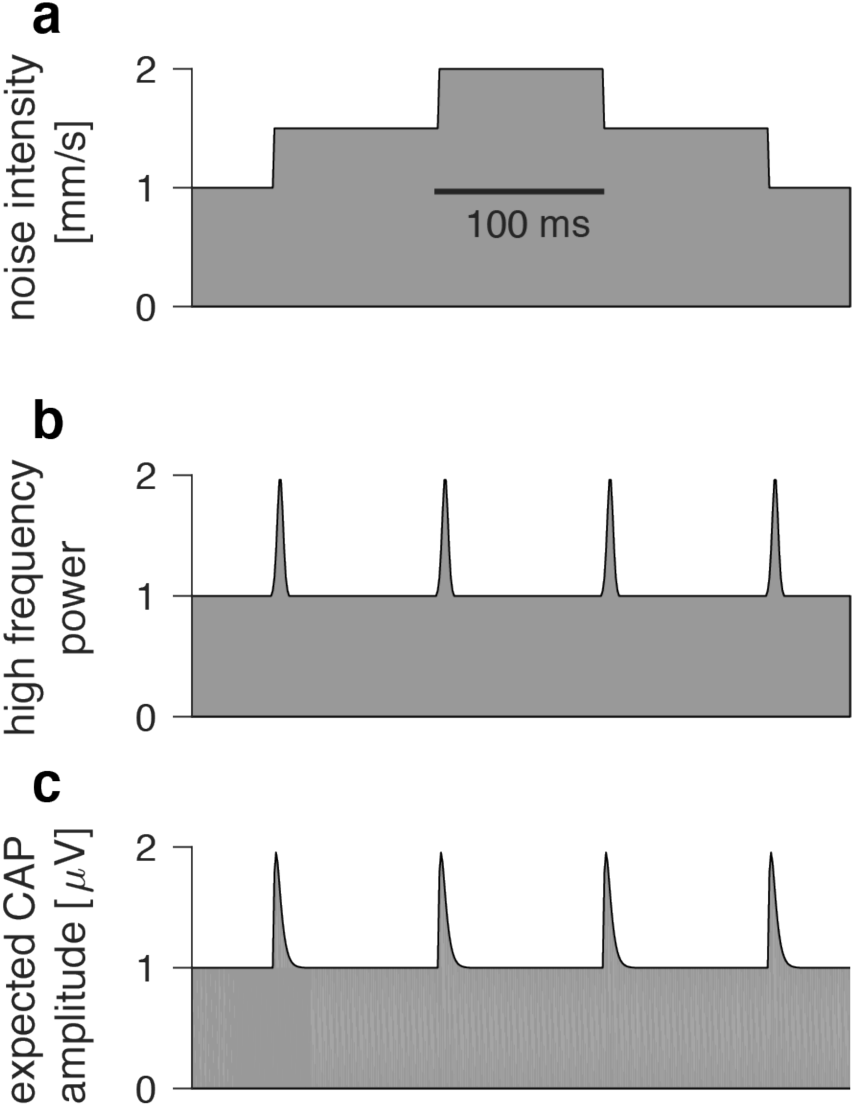
Frequency range fractionation does not explain adaptation dynamics. While our noise stimulus exhibits a relatively constant frequency spectrum, steps in noise intensity (a) can introduce transient increases in high frequency power (b, schematic) and produce transients in the CAP (c, schematic) by briefly recruiting JON that are tuned for higher frequencies. Known tuning profiles of JON are not narrow enough to produce the strong transient that we see in the CAP ^2^ Moreover, the resulting pattern of transients is not consistent with the data (see Fig. 1c), since transients would always positive independent of the direction of the intensity step. We conclude that heterogeneity in frequency tuning cannot explain the CAP dynamics.

**Supplementary Figure 7:**
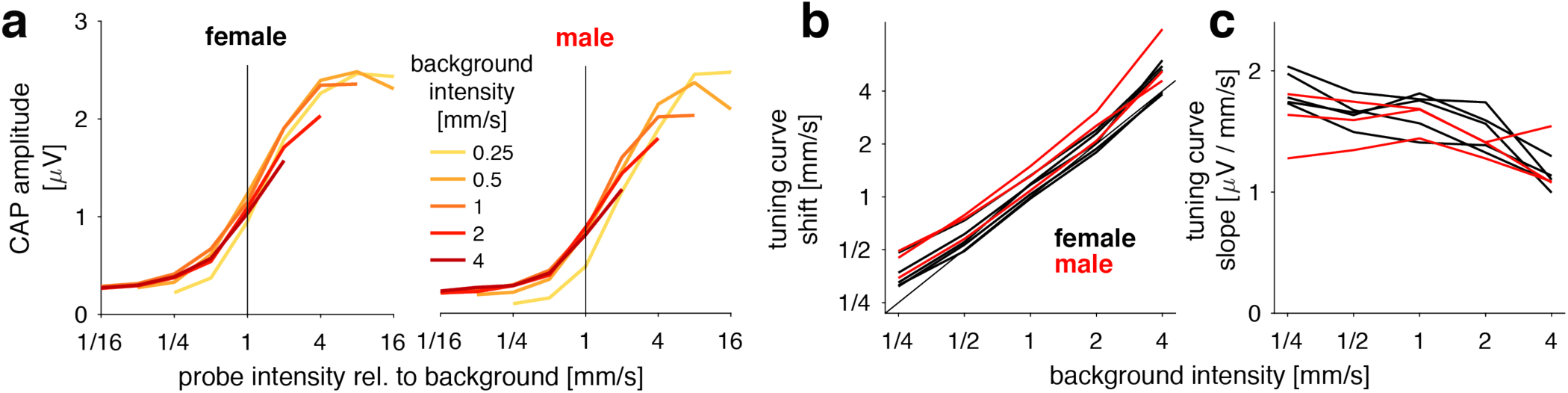
Intensity adaptation is not sex-specific. **a** Intensity adaptation is similar in females (left) and males (right). Representative intensity tuning curves for a single fly (averaged over 20 trials) on an intensity scale relative the background intensity (female data reproduced from Fig. 2) **b, c** The tuning curve shift (b and slope (c) extracted from sigmoidal fits to the set of tuning curves for different backgrounds for all male and female flies (N=5 females (black), N=3 males (red)). Changes in tuning curve parameters with adaptation are similar for males and females, indicating that adaptation is not sex-specific.

**Supplementary Figure 8:**
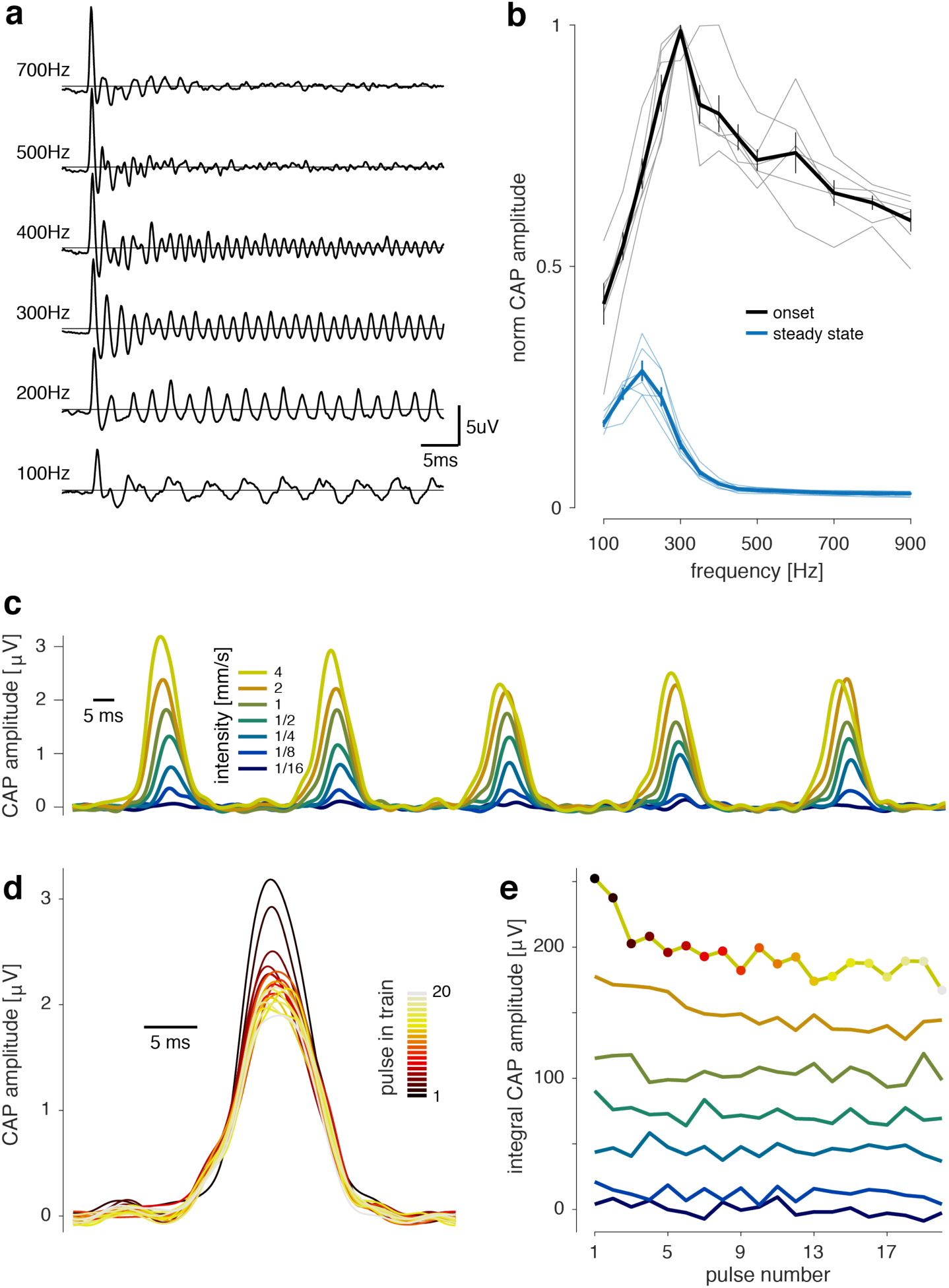
Intensity adaptation for song. **a** Pure tones of different frequencies all elicit adaptation in the CAP (data shown for one fly, averaged over 20 trials, sound intensity of 2 mm s^-1^). **b** Onset (black) and steady-state (blue) responses relative to stimulus frequency (thick lines with error bars are mean±std for N=5 flies, 20 trials; thin lines show tuning for individual flies). The onset and steady-state tuning curve for each fly was normalized such that the maximal onset response was 1.0. Onset responses are given by the average CAP amplitude over the first 10 ms of the stimulus. Steady state responses correspond to the CAP amplitude averaged over 100 ms starting 300 ms after stimulus onset. **c** CAP amplitude for pulse trains (interval 36 ms) at different intensities (color coded, see legend) for one fly (20 trials). **d** Responses to pulses 1 - 20 in a train delivered at 4 mm s^-1^ (same data as in c). Pulse number in train is color coded (see legend). **e** CAP amplitude as a function of pulse number for different pulse intensities (color coded, see legend in c). Colored dots correspond to the pulses shown in d.

**Supplementary Figure 9:**
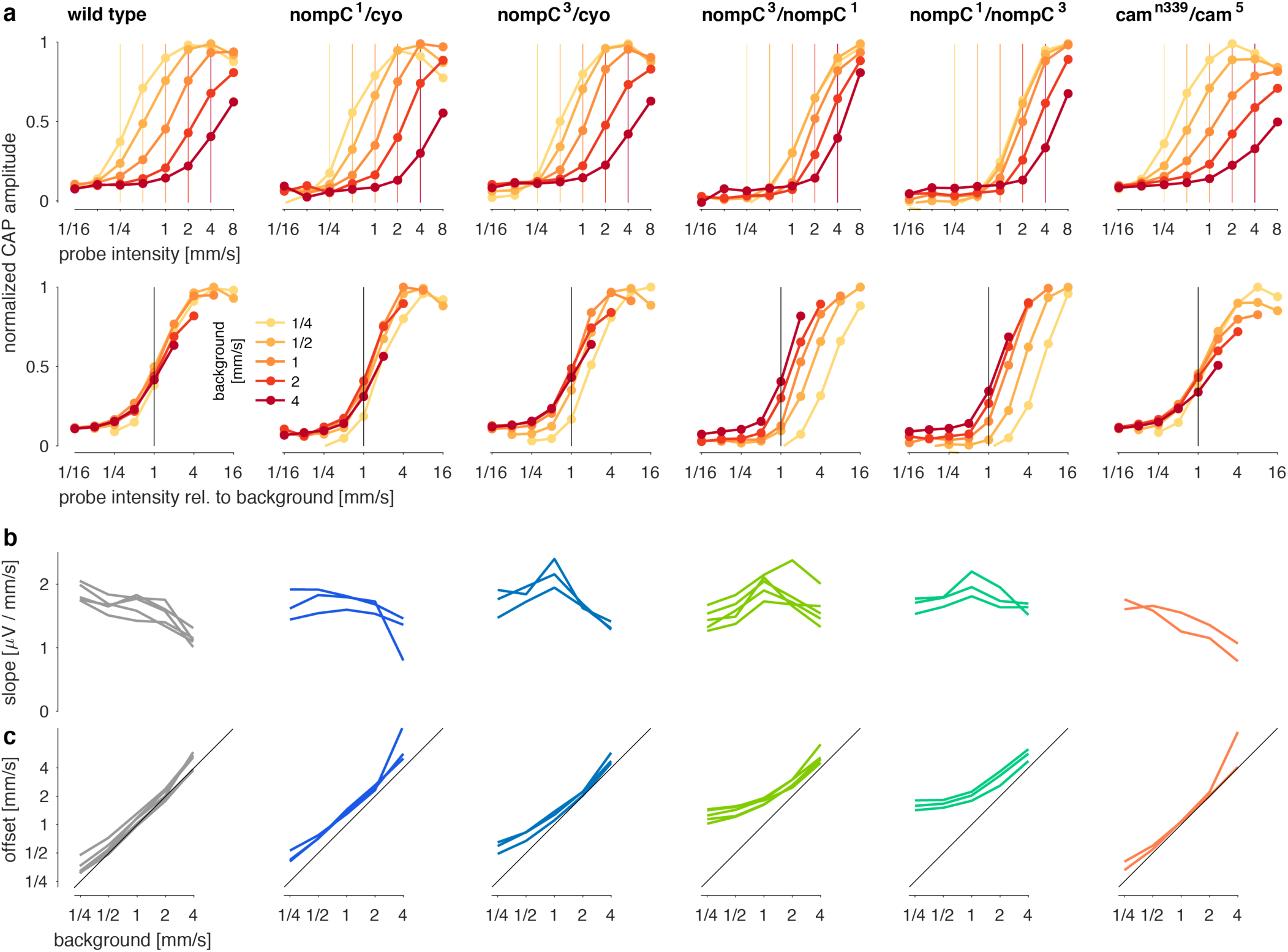
Intensity adaptation persists in nompC and CaM mutants. **a** Intensity tuning curves for different adaptation backgrounds (color coded, see legend) on an absolute (top) and a relative (bottom) intensity scale (compare with Figs. 2B-C). The colored vertical lines (top) indicate the respective background intensity, which corresponds to the black vertical line at 1.0 (bottom) on the relative scale. Plots show tuning curves for one fly of each genotype averaged over 20 trials. **b, c** Slope (b) and offset (c) extracted from sigmoidal fits to the tuning curves for all flies for each genotype. The diagonal line in c corresponds to a perfect match between tuning curve position and background and implies perfect adaptation. Horizontal curves in c would indicate complete absence of adaptation (no tuning curve shift). Tuning curve slopes (b) cover similar ranges across genotypes. Differences in adaptation (c) between strains can be explained by differences in sound sensitivity: some background intensities evoke no responses, and hence no adaptation. The *nompC* heterozygotes (*nompC^1^/cyo* and *nompC^3^/cyo*) exhibit only weakly reduced sensitivity and have largely wild type-like adaptation. In contrast, the *nompC* double mutants (*nompC^3^/nompC^1^* and *nompC^1^/nompC^3^*) are much less sensitive to sound and their adaptation deviates most strongly from wild type - nonetheless adaptation is still observed: for background intensities that evoke strong responses (2 and 4 mm s^-1^), adaptation is like that of wild type. For soft backgrounds, tuning curves still shift with background intensity (a) but adaptation is less complete, as indicated by the deviation of the curves from the diagonal line. Wild type (CS Tully) N=5, *nompC^1^/cyo* N=3, *nompC^3^/cyo* N=3, *nompC^3^/nompC^1^* N=5, *nompC^1^/nompC^3^* N=3, *cam^n339^/cam^5^* N=2.

**Supplementary Figure 10:**
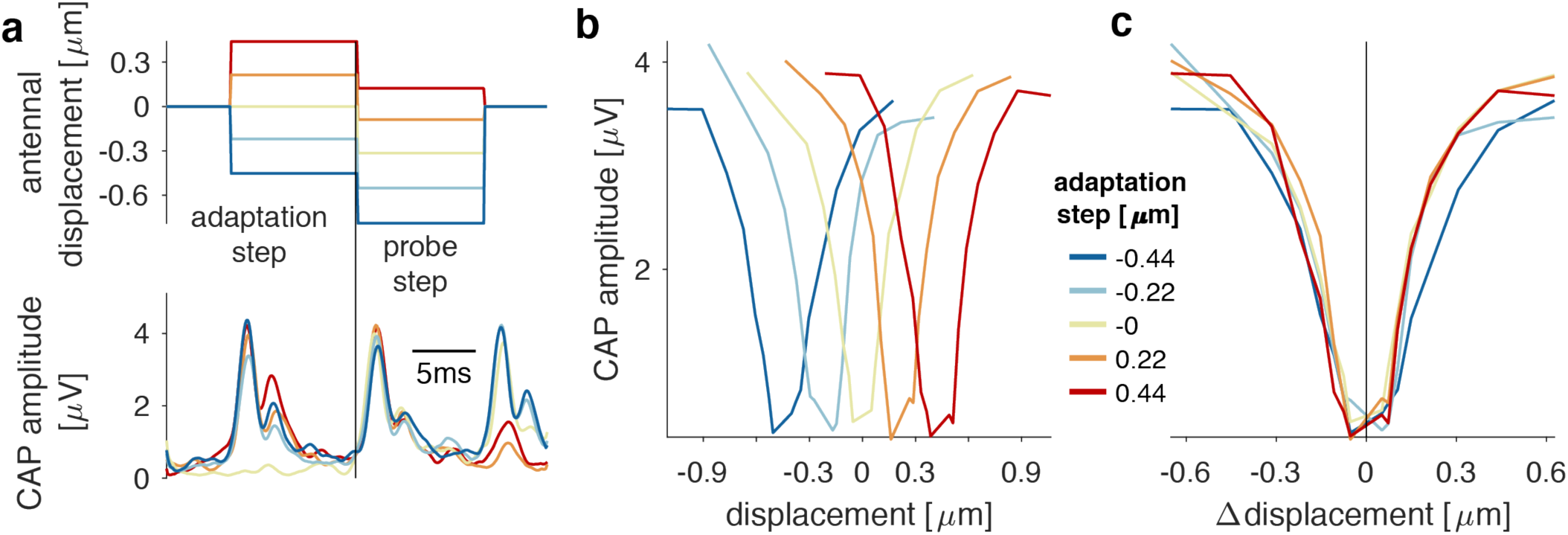
Piezoelectric actuation reproduces offset adaptation. **a** Stimuli for probing offset (mean) adaptation – a piezoelectric actuator was used to step-deflect the antenna to induce offset adaptation (top). Sensitivity to subsequent steps was estimated by measuring the CAP amplitude (bottom; color coded – see legend insert in b). **b** Step tuning curves for different levels of mean adaptation (color coded, see legend). Tuning curve values correspond to the CAP amplitude of the first response peak after the probe step (a) **c** Same curves as in b, with the adaptation background subtracted from the displacement values. All curves overlap, suggesting that JONs encode displacement relative to an additive offset (mean). All plots show responses for one representative fly, averaged over 40 trials.

**Supplementary Figure 11:**
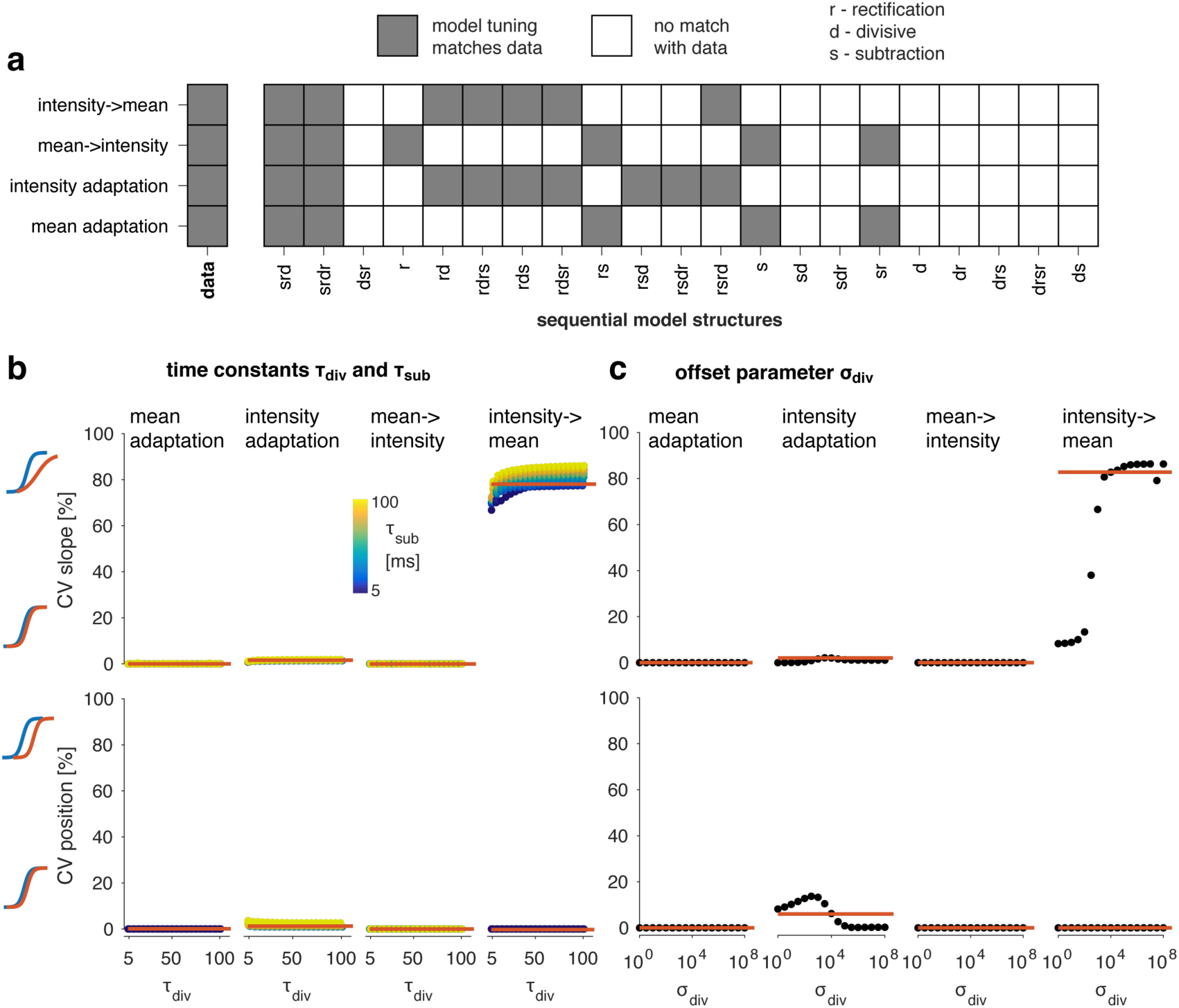
A phenomenological model of mean and variance adaptation. **a** Match between model and data for all models tested. Rows correspond to different adaptation paradigms: The impact of intensity adaptation on step responses (first row), the impact of mean adaptation on intensity tuning (second row), the impact of intensity adaptation or of mean adaptation (3^rd^ and 4^th^ rows). A gray box indicates a match, a white box a mismatch, between model and data. Only two computational motifs (the first two columns) can reproduce our data – both correspond to a sequence of subtractive adaptation (S), rectification (R) and divisive adaptation (D). SRD and SRDR are equivalent – the final rectification step in SRDR is redundant and does not affect model output. **b, c** Model behavior is robust to changes in the time constants for the subtractive and divisive adaptation stage τ_sub_ and τ_div_ (**b**, σ^div^=10^-4^) or the adaptation strength σ_div_ of the divisive adaptation stage (**c**, τ_sub_=30 ms, τ_div_=50 ms). For **b**, τ_div_ are plotted as x-values, while τ_sub_ is color-coded (see legend). Since model behavior is relatively independent of τ_sub_, points for different values of τ_sub_ overlap. Columns in **b** and **c** correspond to different adaptation paradigms: 1) mean adaptation, 2) intensity adaptation, 3) the effect of mean adaptation on intensity tuning, and 4) the effect of intensity adaptation on step responses. Model behavior was quantified using the coefficient of variation (CV) across different adaptation states (e.g. adaptation tone intensity or adaptation step size) for tuning curve slope (top) and position (bottom) extracted from sigmoidal fits (see Methods for details) – pictograms to the left of **b** illustrate tuning curves corresponding to the CVs for a given parameter. The red line corresponds to the CV values obtained for the reference model shown in Fig. 6 that matches our JO data. Most parameter combinations tested produced adaptation like that of the reference model. For small values of the adaptation strength σ_div_ adaptation is too weak to affect tuning curve slope. The model behavior is thus a general property of the computational motif, not the outcome of the specific parameters chosen.

**Supplementary Figure 12:**
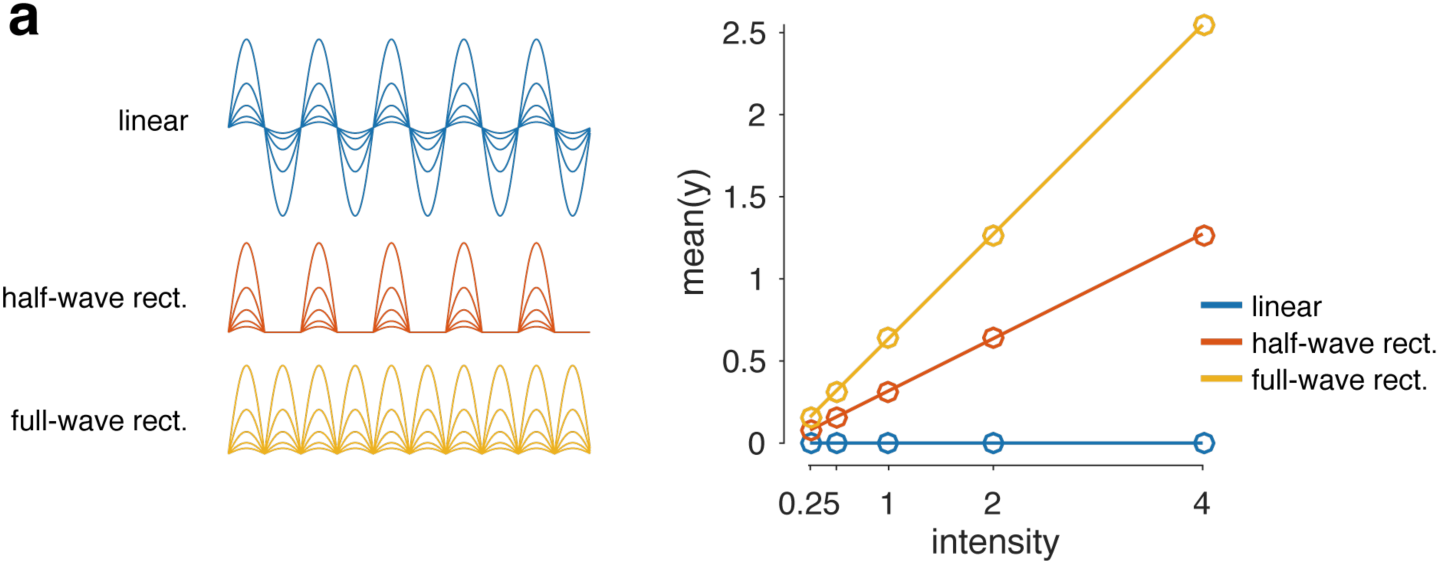
Half-wave and full-wave rectification encode stimulus intensity. **a** A sinusoidal signal of different intensities (left, blue), and the outcome of half-wave rectification (red) or full-wave rectification (orange). The graph shows the mean of the three signals. The mean of the original sinusoidal is zero due to the symmetry of the signal. Full-wave or half-wave rectification lead to the signal mean encoding input magnitude (sound intensity), which is a pre-requisite for variance adaptation in the model.

